# Implantable photonic neural probes for light-sheet fluorescence brain imaging

**DOI:** 10.1101/2020.09.30.317214

**Authors:** Wesley D. Sacher, Fu-Der Chen, Homeira Moradi-Chameh, Xianshu Luo, Anton Fomenko, Prajay Shah, Thomas Lordello, Xinyu Liu, Ilan Felts Almog, John N. Straguzzi, Trevor M. Fowler, Youngho Jung, Ting Hu, Junho Jeong, Andres M. Lozano, Patrick Guo-Qiang Lo, Taufik A. Valiante, Laurent C. Moreaux, Joyce K. S. Poon, Michael L. Roukes

**Affiliations:** Division of Physics, Mathematics, and Astronomy, California Institute of Technology, Pasadena, California 91125, USA; Kavli Nanoscience Institute, California Institute of Technology, Pasadena, California 91125, USA; Department of Electrical and Computer Engineering, University of Toronto, 10 King’s College Rd., Toronto, Ontario M5S 3G4, Canada; Max Planck Institute of Microstructure Physics, Weinberg 2, 06120, Halle, Germany; Krembil Research Institute, Division of Clinical and Computational Neuroscience, University Health Network, Toronto, Ontario, Canada; Advanced Micro Foundry Pte Ltd, 11 Science Park Road, Singapore Science Park II, 117685, Singapore; Institute of Microelectronics, Agency for Science Technology and Research (A*STAR), 2 Fusionopolis Way, #08-02, Innovis, 138634, Singapore; Division of Neurosurgery, Department of Surgery, Toronto Western Hospital, University of Toronto, Toronto, Ontario, Canada; Institute of Biomaterials and Biomedical Engineering, University of Toronto, Toronto, Ontario, Canada

**Author notes:** Equal contribution (Fu-Der Chen, Homeira Moradi-Chameh, Xianshu Luo).

**Keywords:** Neurophotonics, integrated optics, functional imaging, microscopy, biophotonics

## Abstract

**Significance:** Light-sheet fluorescence microscopy is a powerful technique for high-speed volumetric functional imaging. However, in typical light-sheet microscopes, the illumination and collection optics impose significant constraints upon the imaging of non-transparent brain tissues. Here, we demonstrate that these constraints can be surmounted using a new class of implantable *photonic neural probes*.

**Aim:** Mass manufacturable, silicon-based light-sheet photonic neural probes can generate planar patterned illumination at arbitrary depths in brain tissues without any additional micro-optic components.

**Approach:** We develop implantable photonic neural probes that generate light sheets in tissue. The probes were fabricated in a photonics foundry on 200 mm diameter silicon wafers. The light sheets were characterized in fluorescein and in free space. The probe-enabled imaging approach was tested in fixed and *in vitro* mouse brain tissues. Imaging tests were also performed using fluorescent beads suspended in agarose.

**Results:** The probes had 5 to 10 addressable sheets and average sheet thicknesses < 16 μm for propagation distances up to 300 μm in free space. Imaging areas were as large as ≈ 240 μm × 490 μm in brain tissue. Image contrast was enhanced relative to epifluorescence microscopy.

**Conclusions:** The neural probes can lead to new variants of light-sheet fluorescence microscopy for deep brain imaging and experiments in freely-moving animals.

## I. INTRODUCTION

New methods in optogenetics [1–3] and, especially, the advent of fluorescent reporters of neuronal activity, have opened many novel approaches for actuating and recording neural activity *en masse*, through the use of powerful *free-space* single-photon and multi-photon microscopy methods [4–8]. However, existing approaches to functional imaging of the brain have significant limitations. Single-photon (1P) epifluorescence imaging readily lends itself to high frame-rate wide-field microscopy, but, in its simplest implementations, image contrast is hampered by out-of-focus background fluorescence, and the depth of imaging is restricted by the optical attenuation in the tissue. Confocal imaging improves the contrast by optical sectioning, and out-of-focus light is rejected using a pinhole; however, a laser beam must be scanned across each point of the tissue and this significantly slows the image acquisition rate [9]. Multiphoton microscopy is also inherently a point or line scanning method, but because it uses infrared excitation (which provides a longer optical attenuation length [5]), the imaging depth in brain tissue can be extended to ~1 mm and the focus of the light beam can be rastered in three-dimensions to achieve volumetric imaging [5, 10–12].

Light-sheet fluorescence microscopy (LSFM), which is also known as selective-plane illumination microscopy, combines the benefits of fast wide-field imaging, volumetric imaging, and optical sectioning [13]. In conventional LSFM, a thin sheet of excitation light is generated either by cylindrical focusing elements or digitally scanning a Gaussian or Bessel beam [14–16]. The sheet is translated in one dimension across the sample; the fluorescence images are then sequentially collected in the direction perpendicular to the illumination plane to form a volumetric image [17]. With digitally scanned two-photon (2P) LSFM, it is also possible to increase the optical penetration depth [16]. Non-digitally scanned 1P-LSFM is inherently faster than point- or line-scan methods; and since the illumination is restricted to a plane, photobleaching, phototoxicity, and out-of-focus background fluorescence are reduced compared to epifluorescence microscopy. However, conventional LSFM requires two orthogonal objective lenses, and appropriately positioning these largely limits the imaging modality to quasi-transparent organisms (*e.g., C. elegans, Drosophila* embryos, larval zebrafish), chemically cleared mammalian brains [17], and brain slices [18]. An LSFM variant called swept confocally-aligned planar excitation (SCAPE) microscopy, which requires only a single objective, removes these constraints [6, 19]. While *in vivo* calcium neural imaging has been demonstrated using SCAPE in mice [6], miniaturization of the system to be compatible with freely moving animal experiments remains challenging due to the additional optics required.

To make LSFM compatible with non-transparent tissues such as mammalian brains and, eventually, behavioral experiments with freely moving animals necessitates drastic miniaturization of the light-sheet generation and fluorescence imaging compared to today’s archetypical table-top systems. The feasibility of fluorescence microscopy in small and lightweight form factors has already been established by way of head-mounted microscopes for 1P and 2P calcium imaging in mice [4, 20–23], though the endoscopic implantation of the requisite gradient index (GRIN) lenses, with typical diameters of 0.5 - 2 mm, displaces a significant amount of brain tissue.

On the other hand, it remains a formidable and unsolved challenge to generate light sheets by implantable elements at arbitrary brain depths, while minimizing tissue displacement and remaining compatible with a sheet-normal imaging system. For example, in [24], to generate a light sheet perpendicular to the imaging GRIN lens required implantation of a millimeter-scale prism coupled to a second GRIN lens. In another example, in [25], a single light sheet was produced from a microchip using a grating coupler, a glass spacer block, and a metallic slit lens. The overall device was > 100 *μ*m thick and > 600 *μ*m wide, which would displace a significant amount of tissue upon implantation.

Here, we solve these challenges by using wafer-scale nanophotonic technology to realize implantable, silicon-based, light-sheet photonic neural probes that require *no additional* micro-optics. They are fully compatible with free space fluorescence imaging (light collection) outside the brain, where the axis of collection is oriented perpendicular to the light sheets. These silicon (Si) probes synthesize light sheets in tissue using sets of nanophotonic grating couplers (GCs) integrated onto thin, implantable, 3 mm long Si shanks with 50 - 92 *μ*m thickness, widths that taper from 82 - 60 *μ*m along their length, and sharp tips at the distal ends. These prototype photonic neural probes (Fig. 1) are capable of generating and sequentially addressing up to 5 illumination planes with a pitch of ≈ 70 *μ*m. Additionally, the form factor and illumination geometry of the probes open an avenue toward their integration with GRIN lens endoscopes and miniature microscopes, as shown conceptually in Fig. 2(b); offering a singular pathway to rapid, optically sectioned functional imaging at arbitrary depths in the brain.

**FIG. 1:**
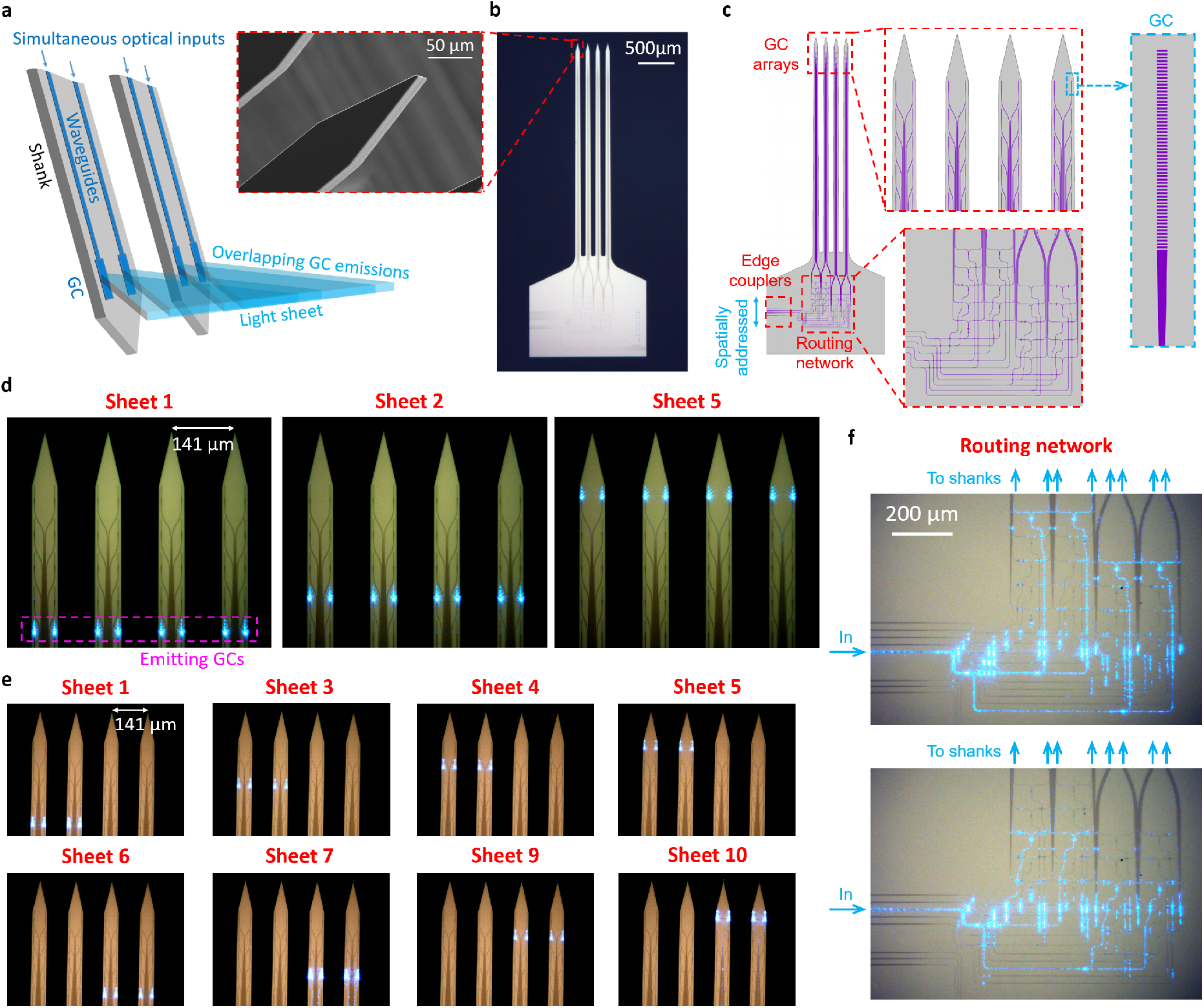
Light-sheet photonic neural probes. (a) Illustration of the light-sheet synthesis method (adapted from [31]). A series of simultaneously fed optical waveguides emits light via a row of grating couplers (GCs) designed for large divergences along the sheet-axis and small divergences along the GC-axis. (b) Optical micrograph of a fabricated neural probe, (inset) scanning electron micrograph (SEM) of the tip of a shank. (c) Top-down schematics of the neural probe. (d-e) Annotated optical micrographs of 2 neural probes with various GC rows emitting light sheets. (d) Neural probe design with sheets generated from 4 shanks. (e) Probe design with sheets generated from 2 shanks (“half-sheet design”). (f) Optical micrographs showing the routing network from the probe in (d) guiding light for optical inputs to 2 different edge couplers. (d-e) have been contrast and brightness adjusted to enhance the visibility of the waveguides.

**FIG. 2:**
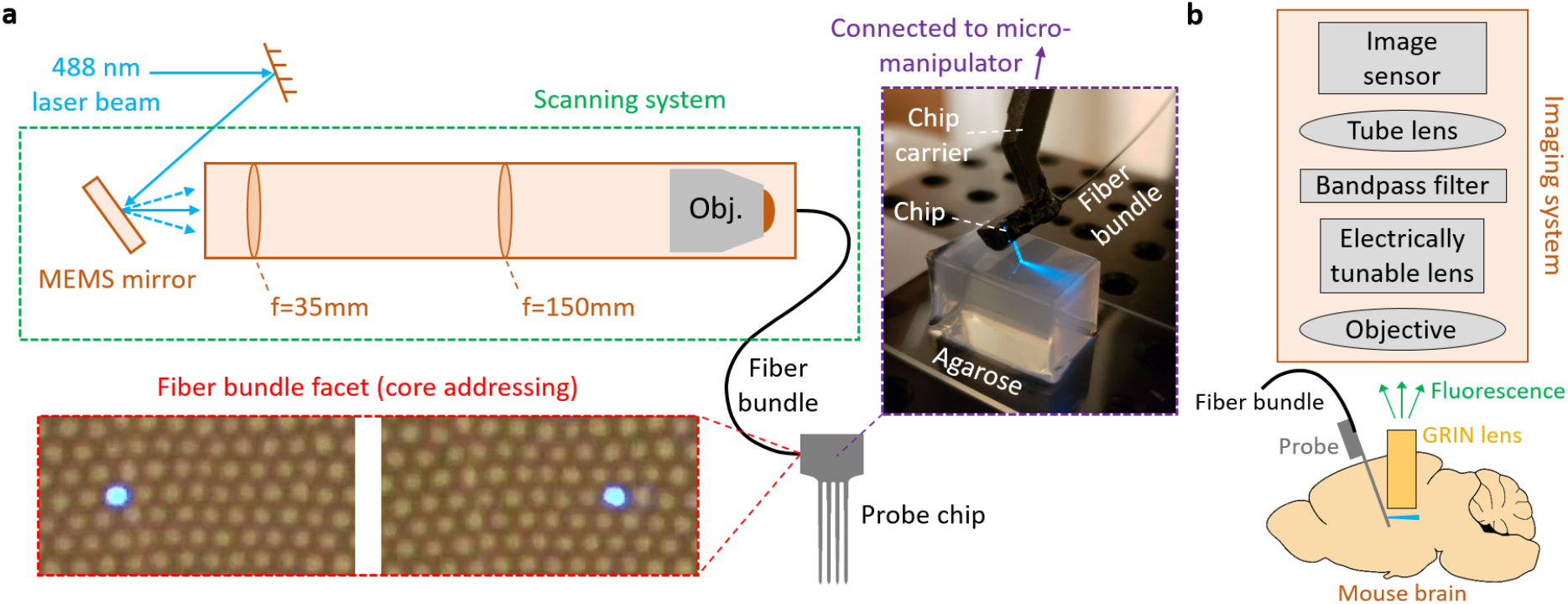
Optical addressing method and proposal for deep-brain photonic-probe-enabled LSFM. (a) Schematic of the optical addressing method (not to scale). The scanning system addresses on-chip edge couplers via spatial addressing of the cores of an image fiber bundle. Bottom inset: micrographs of the distal facet of a fiber bundle connected to the scanning system with different cores addressed (adapted from [31]). Top inset: annotated photograph of a packaged light-sheet neural probe inserted into an agarose block. (b) Illustration of the proposed use of the light-sheet neural probe with a GRIN lens endoscope for deep brain LSFM (not to scale). In this first investigation of the probe functionality, the configuration in (b) has not been demonstrated, and instead, the results here focus on a simpler imaging configuration where lightsheet probe illuminated samples are directly imaged with a fluorescence microscope without a GRIN lens (see Results).

The probes were fabricated on 200 mm Si wafers in a Si photonics foundry for manufacturing scalability and mass-producibility. Elsewhere, we have used this technology to realize photonic neural probes that emit dynamically-reconfigurable, patterned light with cellular-scale beam widths [26] and steerable beams without moving parts [27], adding to a growing number of photonic neural probe demonstrations with increasing levels of integration and sophistication [28–30]. In this work, we employ this integrated nanophotonics technology to realize implantable, microscale probes that form light sheets for imaging over areas as large as ≈ 240 *μ*m × 490 *μ*m in brain tissue. Our preliminary results were reported in [31]. Here, we report in detail the imaging properties of the light-sheet neural probes – characterizing their performance by means of suspended fluorescent beads in phantoms as well as in adult mouse brain slices. A first demonstration of *in vivo* calcium imaging is also reported in the Supplementary Materials.

## II. RESULTS

### A. Photonic neural probes on 200 mm silicon wafers

To ensure that fabrication of our photonic neural probes can be scaled up for dissemination to the neuroscience community, we have adapted from the outset foundry Si photonics manufacturing processes. The neural probes described herein were fabricated in a 200 mm Si photonic line; silicon nitride (SiN) waveguides (135 nm nominal thickness) with SiO_2_ cladding were patterned onto Si wafers, deep trenches were etched in the wafers to define the probe shapes, and the wafers were thinned to thicknesses of 50 - 92 *μ*m. The shank thickness can be reduced in future iterations to 18 *μ*m, as in [26, 31]. The fabrication is more fully detailed in Methods.

The light-sheet neural probe design is shown in Figs. 1(a)-(c). Light is coupled onto the probe chip using fiber-to-chip edge couplers that taper from 5.2 *μ*m in width at the chip facet to single-mode waveguides with widths of 270 – 330 nm. The waveguide-coupled optical power is divided between four to eight waveguides using a routing network consisting of 1 × 2 multimode interference (MMI) splitters [32] and in-plane waveguide crossings [33]. The light is then guided along the implantable shanks via 1 *μ*m wide, multimode waveguides, and subsequently emitted near the distal end of the probe by a row of GCs. Light sheets are synthesized by overlapping the emission from an array of simultaneously-fed GCs. Each row of GCs generates a separate light sheet. The width, period, and duty cycle of the GCs are designed to achieve a large output divergence angle along the width-axis of the sheet, and only a small divergence along the thickness-axis. Nominal lateral GC widths, periods, and duty cycles are 1.5 μm, 440 to 480 nm, and 50%, respectively.

The waveguide routing network is detailed in Fig. S1 in the Supplementary Materials. The photonic components were designed for a wavelength of 488 nm to enable excitation of common fluorophores such as green fluorescent protein (GFP) and green calcium dyes; however, these components can also be designed for green, yellow, and red wavelengths, as we show in [34] for excitation of other fluorophores. The probe shanks are 3 mm in length and separated with a 141 μm pitch; the rows of GCs integrated onto the shanks, each row corresponding to a different sheet, are separated by a 75 μm pitch along the shanks. The shanks taper in width from 82 to 60 μm over their length and each converges to a sharp tip at its distal end.

To rapidly switch between different sheets, we used a spatial addressing approach similar to [35] and as illustrated in Fig. 2(a). An image fiber bundle was epoxied to the probe chip on a common carrier, with each edge coupler on the probe aligned to a different core of the fiber bundle. By actuating the microelectromechanical systems (MEMS) mirror, light was input to a selected core of the fiber bundle and the corresponding input waveguide for a light sheet. The light-sheet switching speed was limited to ≈ 5 ms (0.2 kHz) in the following demonstrations, a constraint arising from the MEMS mirror. Future designs will employ optimized MEMS mirrors operating in resonance mode that can yield switching frequencies > 30 kHz [36]. Videos S1 and S2 demonstrate rapid switching between different light sheets from packaged probes. The fiber bundle used in these first experiments did not maintain polarization, whereas the photonic circuitry was polarization dependent. Therefore, in these first probe prototypes, the fiber bundle must be held still during imaging. This limitation can be overcome in future designs with use of polarization-maintaining multi-core fibers.

Table I summarizes three light-sheet photonic neural probes we have carefully evaluated and report upon in this article. Beam profiles for the three probes are characterized. Probe 1 is used for imaging fluorescent beads and fixed tissue, and Probe 2 is used for *in vitro* imaging. In the table, the “emission angle” refers to the angle of the sheet relative to the normal of the shanks. It is noteworthy that the sheets were designed to emit at an angle of ~ 20° in tissue; this permits implanting the probe next to an imaging lens such that the light sheets can be generated beneath the lens parallel to the focal plane.

**TABLE I:**
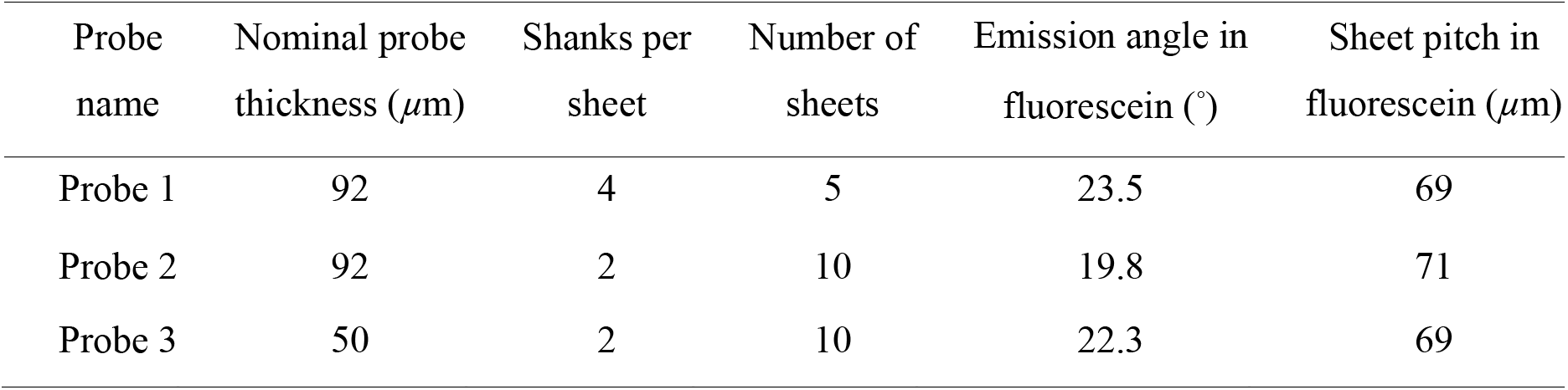
Light-sheet photonic neural probes described in this work

| Probe name | Nominal probe thickness ($\mu\text{m}$) | Shanks per sheet | Number of sheets | Emission angle in fluorescein ($^\circ$) | Sheet pitch in fluorescein ($\mu\text{m}$) |
| --- | --- | --- | --- | --- | --- |
| Probe 1 | 92 | 4 | 5 | 23.5 | 69 |
| Probe 2 | 92 | 2 | 10 | 19.8 | 71 |
| Probe 3 | 50 | 2 | 10 | 22.3 | 69 |

Two probe designs were investigated: a first, in which each light sheet is generated by a row of eight GCs spanning four adjacent shanks (Probe 1, with 5 independent sheets), and a second based on a “half-sheet design”, in which each sheet is generated by a row of four GCs spanning two adjacent shanks (Probes 2 and 3, with 10 independent half-sheets). In principle, the half-sheet design roughly doubles the sheet intensity for a given input optical power to the probe at the expense of a smaller sheet width. More advanced designs can achieve even larger sheet widths by distributing GCs along > 4 shanks at the expense of: 1) a more complex routing network with higher optical losses, and 2) higher input optical powers to the probe chip to achieve a given light-sheet intensity.

### B. Light-sheet generation

The photonic circuitry employed in these devices is designed to provide lower loss for transverse-electric (TE) polarized light. In the following characterization and imaging work, we use TE-polarized optical inputs to the probe chips. The insertion loss of the neural probes (defined here as the ratio of emitted power from the GCs and the input laser power to the scanning system) is summarized in the histograms for Probes 1 and 2 in Fig. S2 in the Supplementary Materials. Probe 3 broke before sheet transmissions were able to be characterized.

Transmission ranged from −38 dB to −20 dB, with a median of about −30 dB. This large variation in transmission was due both to the irregularly positioned individual cores within the fiber bundles and alignment drift during attachment of the fiber to the probe chip. Sheet transmissions measured with a single-mode fiber with optimized alignment typically varied by < 3 dB. In future designs, such transmission variations can be reduced by implementing optimal fiber-to-chip packaging and by employing custom multi-core fibers with a constant core pitch that closely matches that of the on-chip edge couplers. Nonetheless, by modulating the input laser power while switching between sheets or adjusting the MEMS mirror positions for each sheet, these variations can be compensated with the present devices.

We have measured the intrinsic properties of light sheets generated both in free space and in non-scattering fluorescein solutions – characterizing the light-sheet thicknesses, their intensity uniformities, and the magnitudes of associated, higher-order GC diffraction. We determine the in-plane sheet intensity profile by imaging top-down while the probes are immersed in fluorescein solution, Fig. 3(a). When imaging from the side, the sheet thickness is overestimated since out-of-focus light can also be captured. In the free-space method, Fig. 4(a), a coverslip coated on one side with a fluorescent thin film is placed above the probe parallel to the shanks, and a cross-section of the beam profile is imaged on the coverslip. The light sheet intensities were volumetrically profiled versus propagation distance by translating the probe relative to the coverslip.

**FIG. 3:**
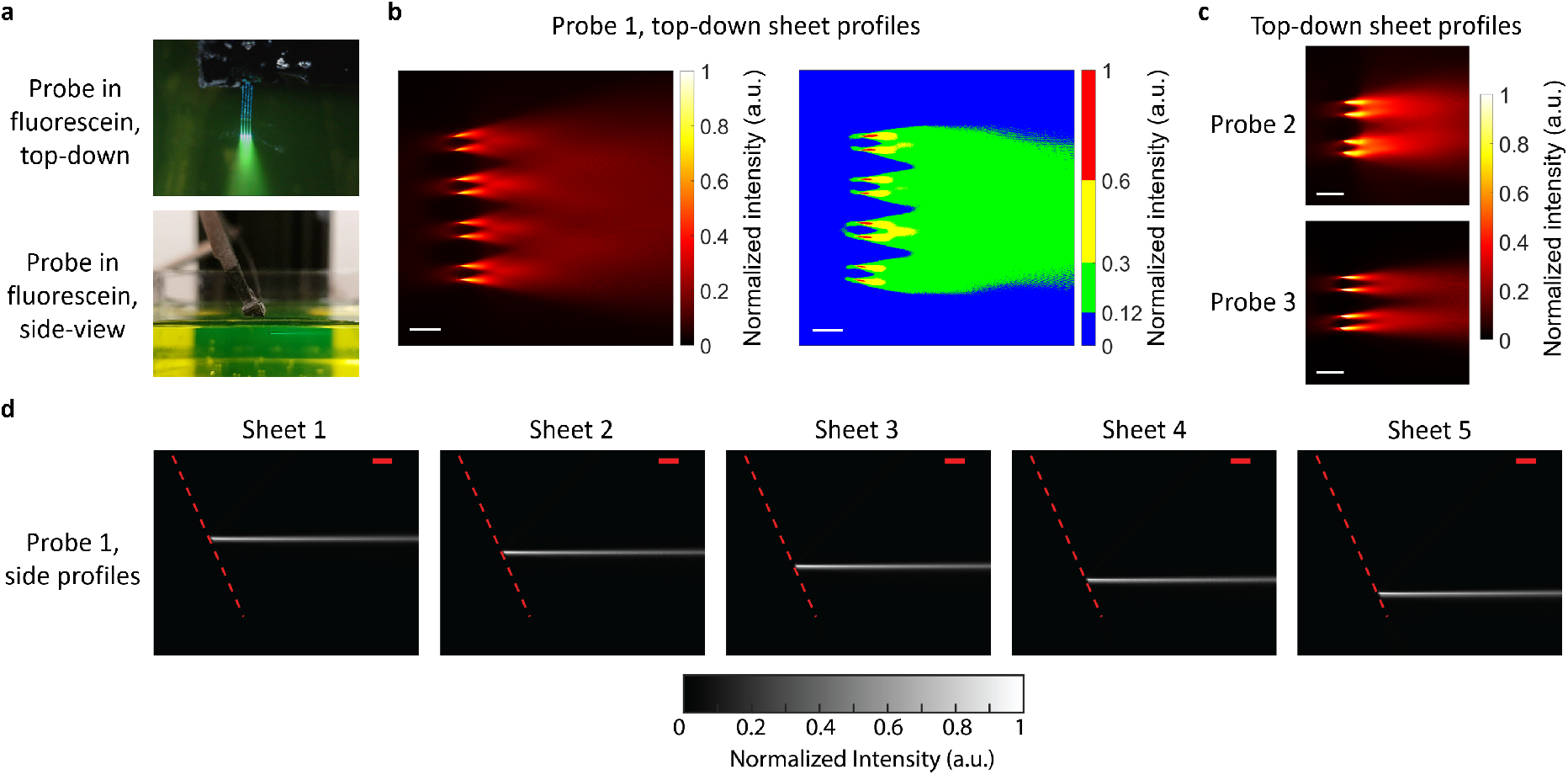
Light-sheet characterization in fluorescein. (a) Top-down and side-view photographs of a light-sheet neural probe immersed in a fluorescein solution. (b) Top-down light-sheet intensity profile for Probe 1 - Sheet 5 imaged with an epifluorescence microscope above the probe. The plot on the right is the sheet profile with a binned color scale to show a semi-uniform sheet region (green) over which the intensity varies by at most 2.5×. (c) Top-down light-sheet intensity profiles for Probe 2 - Sheet 10 and Probe 3 - Sheet 7. (d) Side profile measurements of the light sheets from Probe 1 captured with a second microscope aligned to the side of the fluorescein chamber. The dashed red lines delineate the top surface of the shanks. The scale bars are 100 *μ*m.

**FIG. 4:**
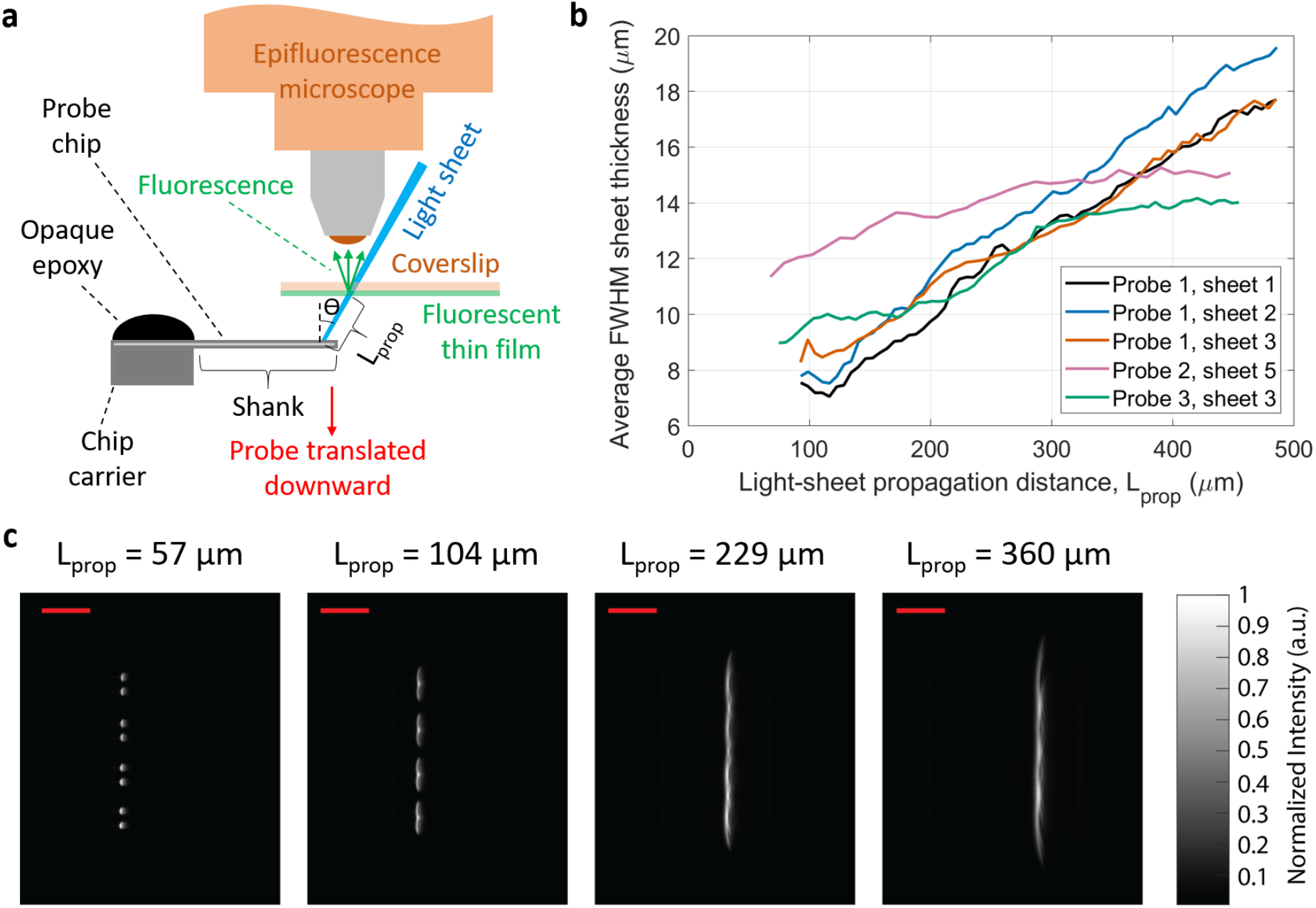
Free-space light-sheet profile measurements. (a) Illustration of the light-sheet profile measurement protocol using a coverslip coated with a fluorescent thin film (not to scale). Fluorescence images of the coverslip provide cross-sectional profiles of the incident light sheet, and vertical translation of the probe enables volumetric profiling. (b) Average full width at half maximum (FWHM) light-sheet thicknesses versus propagation distance for sheets from Probes 1 to 3. The sheet thicknesses are averaged over the sheet width [vertical axis in (c)] for each sheet cross-section. (c) Light-sheet cross-sections imaged at various light-sheet propagation distances, *L_prop_*, for Probe 1. The scale bars are 150 *μ*m.

Figures 3(b)-(c) show top-down fluorescence light-sheet profiles from the probes in fluorescein. The GC emissions diverge and overlap to form regions of moderately uniform illumination. For Probe 1, the semi-uniform region, which we define to be the region where the maximum intensity variations were < 2.5×, is the green region in the binned-color-scale sheet profile of Fig. 3(b). The semi-uniform illumination region forms a continuous sheet at a propagation distance of about 180 μm and spans an area > 0.22 mm^2^. In scattering media such as brain tissue, the semi-uniform illumination region will form at shorter propagation distances away from the probe. The side profiles of the Probe 1 light sheets in fluorescein are shown in Fig. 3(d), and side profiles from Probes 2 and 3 are shown in Fig. S3 in the Supplementary Materials. Weak second-order diffraction results in an additional, upward-pointing beam for each sheet; this is not visible in Fig. 3(d) due to the low second-order diffraction of Probe 1, but it is visible for Probe 2 in Fig. S3. The second-order diffraction profiles were similar to the light sheets, forming “second-order light sheets.” The light sheet optical powers were > 15×, 3×, and 16× larger than the second-order diffraction powers for Probes 1 to 3, respectively.

Figure 4(c) shows light-sheet cross-sections at several propagation distances for Probe 1 imaged with the free-space beam profiling method. The extracted average full width at half maximum (FWHM) light-sheet thickness versus propagation distance for sheets from Probes 1 to 3 are shown in Fig. 4(b). The sheet thicknesses, averaged over the width of each sheet cross-section, are < 16 *μ*m for propagation distances up to 300 *μ*m. Since the coverslip was not perpendicular to the sheet propagation axis, the propagation angle of each sheet is used to convert the thickness of the sheet projection on the coverslip to a sheet thickness corrected for alignment perpendicular to the propagation axis. The apparent reduction of FWHM divergence for Probes 2 and 3 sheets at propagation distances above 300 *μ*m in Fig. 4(b) is a consequence of the evolution of the sheet shape. The full width at 1/*e*^2^ of maximum thickness (Fig. S4 in Supplementary Materials), in general, increases linearly with propagation distance.

Small amplitude fringes are visible in the sheet cross-sections in Fig. 4(c) and the top-down profiles in Fig. 3. These fringes are caused by multi-path interference from the multiple GCs that contribute to each sheet. The interference pattern is related to the differing waveguide path lengths connected to each GC and the coherence length of the laser. In brain tissue, we expect these fringes will be smoothed (suppressed) by the scattering properties of the medium.

### C. Light-sheet fluorescence imaging

We investigate the efficacy of the probes for LSFM by first imaging fluorescent beads suspended in agarose, and then by imaging fixed and *in vitro* brain tissues. Since most miniaturized microscopes today use wide-field 1P fluorescence imaging, we compare the images obtained with the light-sheet probe illumination against those with epi-illumination using the same microscope. Figures 5(a)-(c) illustrate the imaging setup. An electrically tunable lens was attached to the back of the objective to provide fast focus adjustment to the different light-sheet depth planes. When the epi-illumination was on, the input to the probe was off, and vice versa. The comparisons are performed at the same image plane, *i.e*., the tunable lens and microscope objective are not adjusted when switching between light-sheet and epi-illumination. The probe insertion angle was set to orient the light sheets parallel to the top surface of the sample (sheet-normal imaging).

**FIG. 5:**
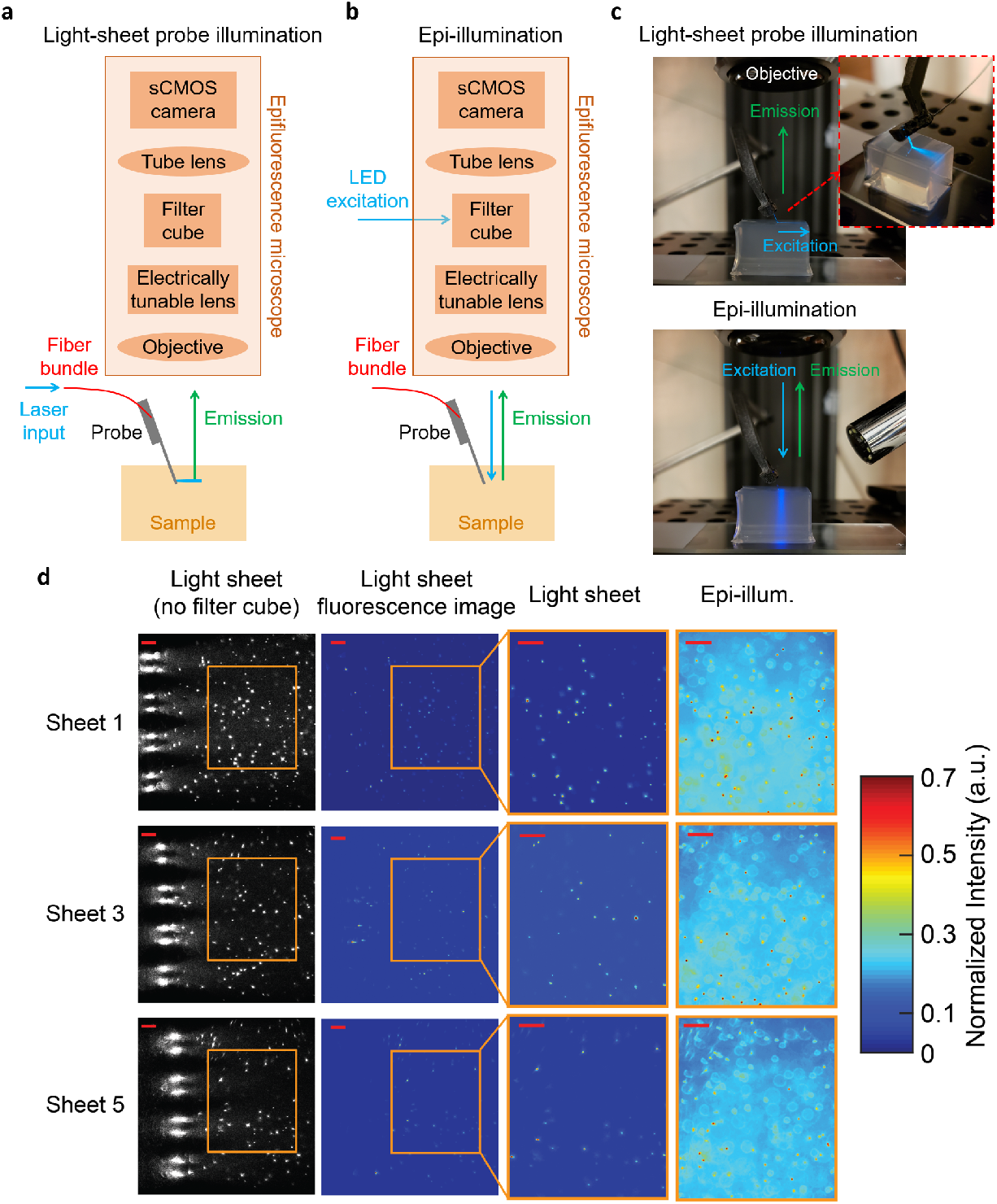
Imaging of fluorescent beads suspended in agarose. (a-b) Illustrations of the imaging apparatus for (a) light-sheet probe illumination, and (b) microscope epi-illumination (not to scale). (c) Photographs of Probe 1 inserted into an agarose block during light-sheet probe illumination and microscope epi-illumination. (d) Imaging of fluorescent beads suspended in an agarose block using light-sheet illumination from Probe 1 (Sheets 1, 3, and 5) and epiillumination. First column: light-sheet illumination images with no filter cube in the microscope path to show both scattered excitation light and fluorescence. The 8 large bright spots at the left of the images are the emitting GCs on the shanks. Second column: fluorescence images with light-sheet illumination and the filter cube in the microscope path. Third and fourth columns: fluorescence images of the regions of interest delineated by the orange boxes with light-sheet and epi-illumination, respectively. The epi-illumination images were captured at the same focal planes as the corresponding light-sheet images. The second to fourth columns are normalized to the maximum intensity in each image and the color scale is truncated at 0.7 to enhance bead visibility. The scale bars are 50 *μ*m.

To demonstrate optical sectioning, Probe 1 was inserted into an agarose block containing 3 *μ*m diameter fluorescent beads. Figure 5(d) shows the fluorescence images captured using 3 of the sheets of Probe 1 compared with epifluorescence images. Significantly out-of-focus beads and fluorescence are not present with light-sheet probe illumination. This yields a dramatic reduction of the background intensity in comparison with epi-illumination. We quantify the gain in contrast in imaging experiments with tissue slices, discussed next. Video S3 shows a simple proof-of-concept volumetric imaging example. The video demonstrates fluorescence imaging of fluorescent beads in an agarose block with switching between three of the probe-generated sheets, and synchronized focus switching enabled by the electrically tunable lens.

We subsequently imaged the hippocampus in a fixed brain slice obtained from a Thy1-GCaMP6s mouse. Figure 6 shows the probe- and epi-illuminated fluorescence images captured following insertion of Probe 1 into the fixed tissue. The tissue was about 1 mm thick, and imaging was performed with Sheets 3 and 4. Again, the probe-illuminated images showed remarkably less background fluorescence than epi-illumination. Neurons are observable over a sheet area of ≈ 240 *μ*m × 490 *μ*m for Sheet 3, and different neurons are visible with Sheet 3 versus Sheet 4 illumination. The neurons in the image from Sheet 4, which was 69 *μ*m deeper in the tissue than Sheet 3, appear less in focus; this is due to the scattering of the fluorescence emission in the tissue.

**FIG. 6:**
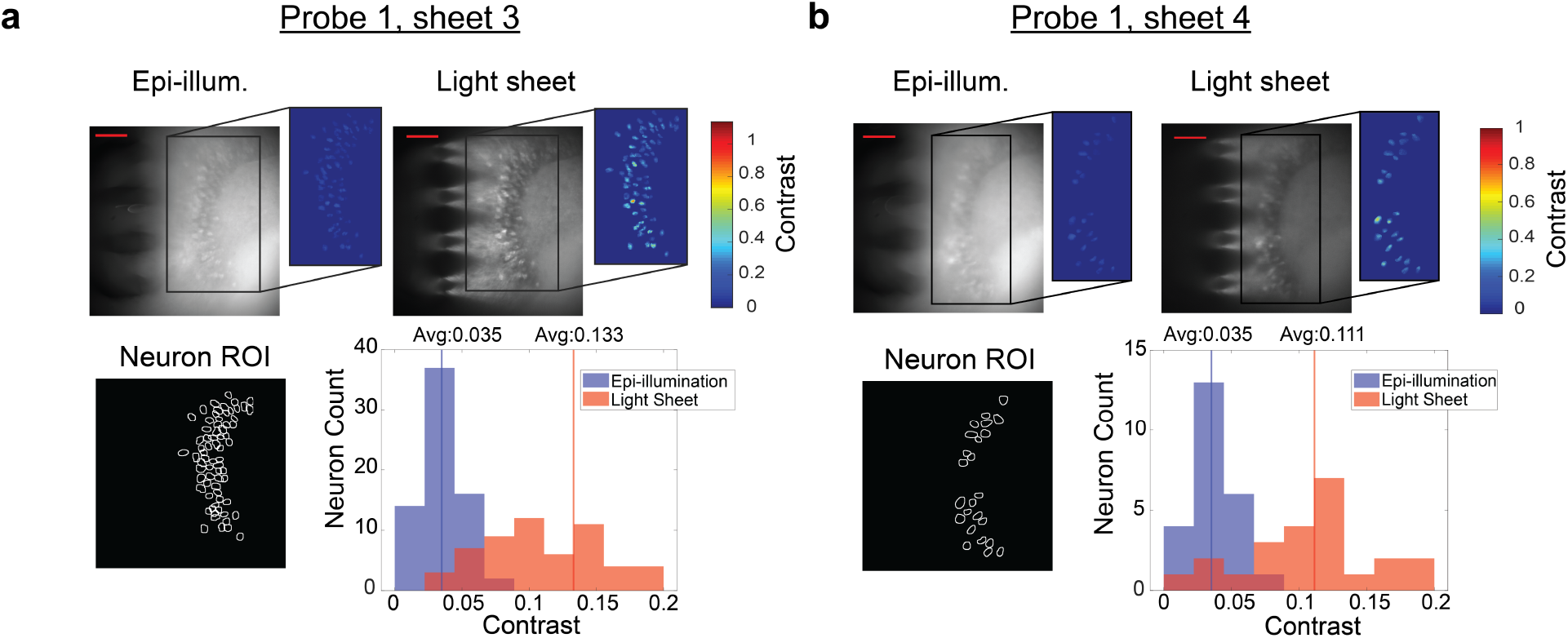
Comparison of light-sheet neural probe illumination and microscope epiillumination for fluorescence imaging of fixed brain tissue (dentate gyrus) from a Thy1-GCaMP6s mouse. Two adjacent light sheets from Probe 1 were used: (a) Sheet 3, and (b), Sheet 4. Sheet 3 was ≈ 60 *μ*m in depth from the surface of the brain tissue, and Sheet 4 was 69 *μ*m deeper than Sheet 3. Top row: fluorescence images for epi- and light-sheet illumination with insets indicating the contrast of neurons within a region of interest of high neuron density. The scale bars are 100 *μ*m. Bottom row: regions of interest (ROIs) of identified neurons and corresponding histograms of image contrast for the identified neurons; the contrast of each neuron is the average over its ROI.

To quantify the difference in contrast between probe- and epi-illumination, an algorithm described in Supplementary Note 1 (see Supplementary Materials) is applied to identify the neurons in each image, and the neurons found in both images are selected (Fig. 6 “Neuron ROI”) for contrast analysis using the definition of contrast in Supplementary Note 1. Figure 6 shows the distributions of the image contrasts of the identified neurons. The contrast distributions of the two illumination methods are statistically different (*p* < 0.001, two-tailed Wilcoxon signed-rank test), with the average contrast for light-sheet illumination higher than that of epi-illumination by 3.8× for Sheet 3 and 3.2× for Sheet 4. 98.6% and 100% of the neurons for Sheets 3 and 4, respectively, exhibit higher contrast using light-sheet illumination compared to epi-illumination. The color insets in Fig. 6 show the contrast of each pixel within each neuron ROI, while the histograms show neuron contrasts that are averaged over each neuron ROI. The illumination intensities for the fixed tissue and *in vitro* imaging are discussed in Supplementary Note 2.

Photonic neural probe tests were also performed for *in vitro* functional calcium imaging using a 450 *μ*m thick brain slice, prominently featuring the auditory cortex, from a Thy1-GCaMP6s mouse. Preparation of the tissue is described in Methods. For increased neuronal activity, the brain slice was perfused with an artificial cerebral spinal fluid (aCSF) solution containing 4-aminopyridine (4-AP) [37]. Figure 7(a) shows maximum projection images over time from the probe- and epi-illumination videos of the auditory cortex region of the brain slice. Sheet 5 from Probe 2 was used, and the probe was inserted into the brain slice such that Sheet 5 was ≈ 60 *μ*m in depth from the surface of the slice. The data analysis procedure for neuron identification and extraction of the fluorescence change, Δ*F/F*, is described in Supplementary Note 3. Figure 7(c) shows the Δ*F/F* time traces of the 16 identified active neurons using probe-illumination, and Δ*F/F* values as large as 5.5 were observed. Figure 7(d) shows the image contrast of 5 of the neurons at the peaks of all observed events; the neurons were selected with the criterion that at least 5 events were recorded for both light-sheet and epi-illumination. Higher image contrast is observed for light-sheet compared to epi-illumination for 4 of the 5 neurons (*p* < 0.01, two-tailed Wilcoxon rank-sum test); a possible explanation for the lower light-sheet contrast of the one neuron is that its depth may have been outside or on the periphery of the sheet. The ratios of the median light-sheet- and epi-illumination neuron contrasts were 6.71, 0.77, 2.04, 2.46, and 3.39. Samples of calcium imaging video with both probe- and epi-illumination are shown in Videos 1 and 2.

**FIG. 7:**
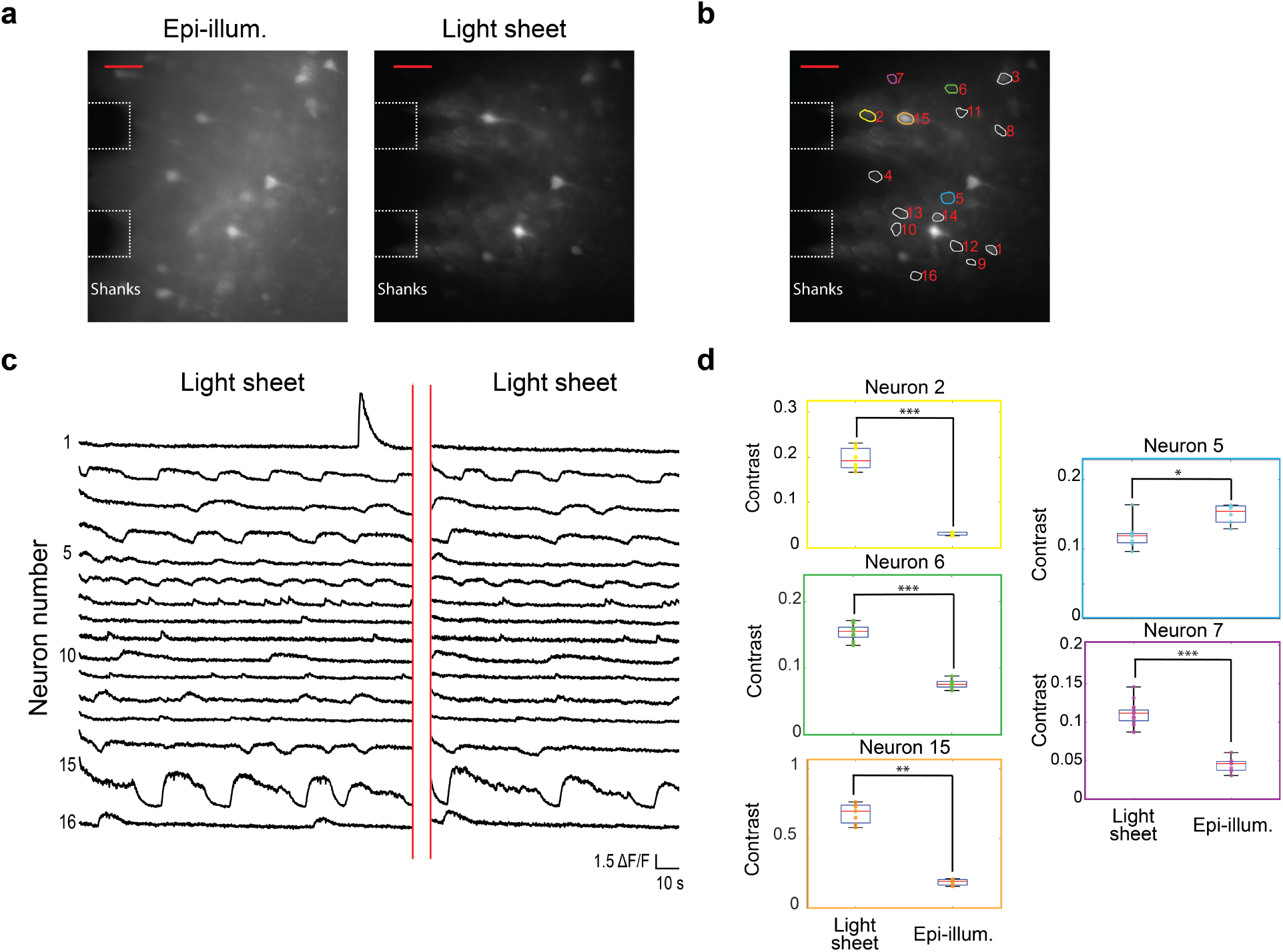
*In vitro* functional calcium imaging of a brain slice from a Thy1-GCaMP6s mouse. (a) Maximum projection images of 142 s and 92 s segments of the recorded video for light-sheet and epi-illumination, respectively, with annotations showing the approximate positions of the shanks in the image plane. The scale bars are 50 *μ*m. (b) Light-sheet maximum projection image with ROIs for identified active neurons shown. (c) Fluorescence change, Δ*F/F*, time traces of all active neurons identified in (b). (d) Box plots showing the image contrast of 5 neurons at the peaks of all events recorded for light-sheet and epi-illumination; the numbers and colors of the box plots correspond to the ROIs in (b). Asterisks indicate significant group differences. * denotes *p* < 0.05, ** denotes *p* < 0.01 and *** denotes *p* < 0.001, two-tailed Wilcoxon rank-sum test. A sample of the calcium imaging video with light-sheet probe illumination is presented (Video 1, 12.9 MB, mp4). A sample of the calcium imaging video with epi-illumination is also presented (Video 2, 29.8 MB, mp4). The videos are accelerated 5×. The Δ*F/F* time traces for epiillumination are shown in Fig. S6 in the Supplementary Materials.

During calcium imaging experiments, illumination was alternated between probe- and epi-illumination. The full time-traces are shown in Fig. S6 in the Supplementary Materials. Due to the larger background fluorescence of epi-illumination, the apparently larger Δ*F/F* for light-sheet compared to epi-illumination does not necessarily represent a larger signal-to-noise ratio of the calcium events. A direct comparison of signal-to-noise ratio for calcium events under these two illumination conditions is beyond the scope of this work.

To investigate the operation of the probes in tissues with a higher density of labeled neurons, tests were also performed on a green calcium dye loaded (Cal-520 AM, AAT Bioquest), 450 *μ*m thick, cerebellum brain slice from a wild type mouse. The tissue preparation is described in Methods, and 4-AP was added to the aCSF perfusion solution. Probe 2 was inserted into the brain slice, and light-sheet illumination was applied from Sheet 10, which was positioned < 50 *μ*m in depth from the brain slice surface (Fig. 8). Full fluorescence time-traces with the illumination cycled between probe- and epi-illumination are shown in Fig. S8 in the Supplementary Materials. Samples of the calcium imaging video are shown in Videos 3 and S4. The labeled cells are likely a combination of neurons and glial cells. The data analysis procedure is described in Supplementary Note 3. For the probe illumination in Fig. 8, Δ*F/F* values as high as 4.3 were observed, and 73 cells were identified. The variation in maximal Δ*F/F* values in Fig. 8(c) may arise from a combination of the position of the cell within the sheet (both laterally and in depth) as well as the magnitude of the calcium events. A complication in this experiment arises from the penetration depth of the dye into the slice during bath-loading; this limits the thickness of labeled tissue available to contribute to background fluorescence during epi-illumination. As a result, the image contrast enhancement of light-sheet versus epi-illumination is expected to be less than our results with Thy1-GCaMP6s mouse brain tissue (Figs. 6 – 7), where the labeling is more uniform in depth. This is confirmed by the minor contrast difference between light-sheet and epi-illumination observed for the Cal-520 AM loaded brain slice (Videos 3 and S4, Fig. S8 in the Supplementary Materials), relative to the significant contrast enhancement of light-sheet illumination in Figs. 6 – 7.

**FIG. 8:**
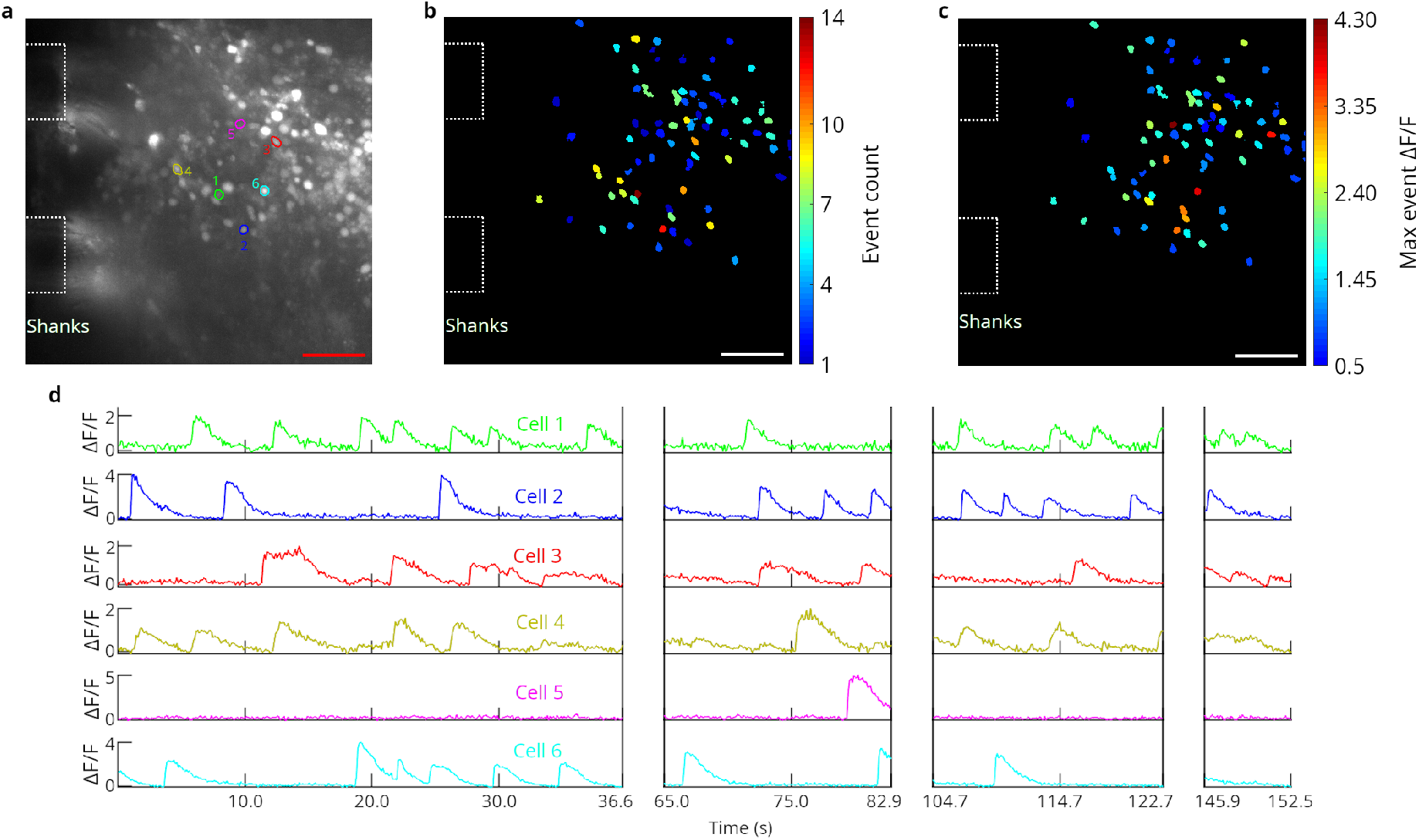
*In vitro* functional calcium imaging of a Cal-520 AM loaded brain slice from a wild type mouse using light-sheet neural probe illumination. (a) Maximum projection fluorescence image of 36.6 s of the recorded video with annotations showing the approximate positions of the shanks in the image plane. (b) Maximum observed Δ*F/F* and (c) number of calcium events observed for all identified cells. (b-c) are of the same scale as (a), and the scale bars are 50 *μ*m. (d) Fluorescence change, Δ*F/F*, time traces of 6 cells; the first 4 cells had the highest number of events and the last 2 had the highest peak Δ*F/F* among the remaining cells. The ROIs for these cells are shown in (a) with colors and numbers corresponding to the time traces in (d). Breaks in the time traces correspond to times when the illumination was switched to epi-illumination. A sample of the calcium imaging video with light-sheet probe illumination is presented (Video 3, 19.6 MB, mp4). The video is real-time.

We have carried out initial *in vivo* tests as shown in Fig. S11 and discussed in Supplementary Note 4 (see Supplementary Materials). For these experiments, a light-sheet probe was inserted approximately < 200 μm deep into the parieto-temporal lobe of an anesthetized Thy1-GCaMP6s mouse at the approximate location of the somatosensory cortex. Time-dependent fluorescence using probe-illumination was observed with a maximum Δ*F/F* of 0.12, and contrast enhancements were observed compared to epi-illumination. In this case, it proved difficult to establish statistical comparisons due to the low number of neurons exhibiting activity in the anesthetized mouse. The probe used for this test was an earlier prototype, which pre-dated our foundry-fabricated probes; details of these earlier devices are listed in Supplementary Note 5.

## III. DISCUSSION AND CONCLUSION

We have conceived of and demonstrated a new paradigm for implantable photonic neural probes that enables lensless delivery of multiple addressable light sheets. These can facilitate 1P-LSFM at arbitrary depths in mammalian brains and other non-transparent tissues. The light-sheet profiles were experimentally characterized, and the probes were validated by fluorescence imaging in fixed tissue and by functional imaging *in vitro*. This imaging approach requires *no active components* on the probe, which can otherwise induce deleterious tissue heating. By contrast, miniaturized forms of digitally scanned 1P- or 2P-LSFM would likely require actuators on the probe or in close proximity thereof. As the light-sheet neural probes are passive, the ultimate volumetric imaging rate is similar to that of conventional light-sheet imaging systems. In our probe-based implementation, it is limited by external components that include the electrically tunable lens, MEMS mirror, and the image sensor. The apparatus we employ here is not yet fully optimized to achieve maximum volumetric imaging rates and is primarily limited by the tunable lens, which has a response time of 25 ms. Other system components are faster; the MEMS mirror step time is ~ 5 ms, and the maximum full frame rate of the camera is 101 frames per second. Accordingly, with optimized component choices and engineering of the imaging system, volumetric imaging rates ≥ 30 volumetric scans per second will be attainable [38]. Although continuous-wave light was used in our experiments, future implementations can employ pulsed light to mitigate any potential phototoxicity and photodamage.

The light sheets created by our probes are synthesized from an incoherent sum of multiple GC optical emissions. We expect sheets generated by neural probes or by conventional light-sheet microscopes to be similarly affected by scattering within brain tissue. Optical scattering has been evaluated for conventional light-sheet microscopes in [39] and for nanophotonic GC emissions in [26, 29].

In future iterations of our probe designs, their photonic circuits can be further optimized by leveraging state-of-the-art integrated photonic technology. For example, the sheet density may be increased by integrating multiple photonic layers [40]. Also, the optical transmission of the probes can be increased by roughly an order of magnitude with efficient fiber-to-chip edge couplers [41] and optimized low-loss components; the fiber-to-chip coupling efficiency of the edge couplers in this work was limited to ≈ 14% with optimal alignment [34]. Optimized packaging solutions can also mitigate transmission variations amongst light sheets and improve the thermal stability of the packaged probes. (The latter may eliminate the turn-on-transient documented in Fig. S10 in the Supplementary Materials.) Optimization of probe transmission and packaging can also minimize potential tissue heating arising from the packaged probes absorbing extraneous scattered light from on-chip photonic circuitry and edge couplers.

With their micro-scale form factors, ultrathin profiles, and their compatibility with sheet-normal imaging using implantable GRIN lens endoscopes, the light-sheet photonic neural probes we have demonstrated herein can engender exciting and powerful new variants of LSFM, both for deep brain imaging and for behavioral experiments with freely-moving animals. Beyond LSFM imaging, these neural probes can also be used for laminar optogenetic neural stimulation, *e.g*., for addressing *individual* cortical layers. When combined with a new class of implantable neural probes containing photodetector arrays that is now emerging [42], they can enable complex image reconstruction realized by means of a complete, implantable lensless imaging system.

## IV. METHODS

### Photonic neural probe fabrication

The neural probes were fabricated on 200 mm diameter Si wafers at Advanced Micro Foundry (AMF). First, the 1.48 *μ*m and 135 nm thick SiO_2_ bottom cladding and SiN waveguide layer were deposited by plasma enhanced chemical vapor deposition (PECVD). Fully etched SiN waveguides were formed using deep ultraviolet (DUV) lithography followed by reactive-ion etching (RIE), and the 1.55 *μ*m thick PECVD SiO_2_ top cladding layer was then deposited. Deep trenches were etched to define the probe shape and form facets for edge couplers. Finally, as in [43], backgrinding was used to thin the wafers to ≈ 50 - 92 *μ*m, which exposed the deep trenches and separated the probes on the grinding tape (autodicing). Chemical mechanical planarization (CMP) was used for layer planarization during the fabrication. The fabrication process and waveguide characteristics are described in more detail in [34].

### Neural probe packaging

The probe chip was first epoxied to a 3D-printed chip carrier. The image fiber bundle (Fujikura FIGH-06-300S) was connected and aligned to the scanning optical system, Fig. 2(a). The fiber bundle was aligned and then UV-epoxied to the probe chip and the chip carrier; the emitted optical power from the probe was monitored during the process. The probe chip (excluding the shanks) and the fiber bundle were then coated with optically opaque epoxy to block stray light not coupled to the on-chip waveguides. The chip carrier had a steel rod attached to the proximal end, and this steel rod was connected to additional rods to mount the packaged probe on a 4-axis micro-manipulator (QUAD, Sutter Instrument Company, Novato, CA, USA).

### Spatial addressing of the neural probes

The 2-axis MEMS mirror in the external scanning optical system, Fig. 2(a), had a nominal maximum mechanical tilt angle of ±5.5° and a mirror diameter of 3.6 mm (A7B2.1-3600AL, Mirrorcle Technologies Inc., Richmond, CA, USA). The scanning system used bi-convex lenses with 35 mm and 150 mm focal lengths and a 20× objective lens (Plan Apochromat, 20 mm working distance, 0.42 numerical aperture, Mitutoyo Corporation, Kawasaki, Japan). The loss of the scanning system (from the input laser beam to the distal facet of the image bundle) was typically 40 - 60%. The 488 nm laser (OBIS 488 nm LS 150 mW, Coherent Inc., Santa Clara, CA, USA) was fiber-coupled to a single-mode fiber (460-HP, Nufern Inc., East Granby, CT, USA), which was connected to a fiber collimator that formed the free-space laser beam input to the scanning system. The input beam was gated by a mechanical shutter. The input polarization to the scanning system was set via an inline fiber polarization controller.

### Fluorescein beam profiling

The neural probes were dipped into 10 *μ*mol fluorescein solutions (pH > 9), Fig. 3(a). Top-down sheet profiles were measured using an epifluorescence microscope above the probe. Side profiles showing the sheet thicknesses were measured using an additional microscope positioned at the side of the chamber containing the fluorescein. One of the walls of the chamber was removed and replaced with a coverslip to create a viewing port with low optical distortion for the side microscope. Bandpass optical filters on both microscopes rejected excitation light from the probe and transmitted the emission light from the fluorescein. The insertion axis of the micro-manipulator holding the probe was angled such that the sheets propagated parallel to the surface of the fluorescein solution with the probe immersed.

### Free-space beam profiling

Coverslips with a fluorescent thin film were fabricated by mixing fluorescein (free acid) powder with SU-8 photoresist [44] and spin coating it onto ≈ 170 *μ*m thick coverslips. After curing, an ≈ 8 *μ*m thick fluorescent thin film was formed on one side of the coverslips, which were then cleaved in half to prevent the edge bead from limiting the probe-to-coverslip distance. The coverslip was fixed above the probe with the shanks, coverslip, and optical table parallel (Fig. 4). The fluorescent film was on the bottom side of the coverslip, closest to the probe. An epifluorescence microscope above the coverslip imaged the fluorescent patterns created by the intersection of the probe light sheets and the thin film. Vertical translation of the probe enabled volumetric profiling of each sheet for measurements of the sheet thickness and propagation angle. The sheet propagation angles were used to convert the micro-manipulator vertical translation step size into sheet propagation distance step sizes, and the angles were also used to calculate the sheet thicknesses from the angled projections on the coverslip. To verify the uniformity and linearity of the thin film’s fluorescence, volumetric profiles were captured over input optical powers spanning roughly an order of magnitude and at multiple positions on the coverslip. Measured average sheet thicknesses varied by < 2 *μ*m throughout the trials.

### Fluorescent beads in agarose

To prepare the agarose blocks with fluorescent beads, 100 mg of agarose powder was mixed with 10 mL of Milli-Q water to form a 1% agarose solution. The solution was heated until boiled, and after cooling, 50 *μ*L of yellow-green fluorescent microbeads (3 *μ*m bead diameter, 2.5% concentration, Magsphere, Pasadena, CA, USA) were mixed into the solution. The solution was placed on a rocker to evenly distribute the beads, and then, poured into a plastic mold and stored in a refrigerator until solidified. The intensity scales of the grayscale images of fluorescent beads in Fig. 5(d) were set with the bottom and top 1% of all pixel intensities saturated.

### Imaging apparatus

The fluorescence imaging apparatus, Figs. 5(a) and (b), includes an epifluorescence microscope (Eclipse FN1, Nikon, Tokyo, Japan) with an sCMOS camera (Zyla 4.2 PLUS, Andor Technology Ltd., Belfast, UK) and an EGFP filter cube (49002, Chroma Technology Corporation, Bellows Falls, VT, USA). A 10× objective lens (Mitutoyo Plan Apochromat, 34 mm working distance, 0.28 numerical aperture) was used for the beam characterization and *in vitro* Cal-520 AM brain slice imaging. A 20× objective (Mitutoyo Plan Apochromat, 20 mm working distance, 0.42 numerical aperture) was used for the fluorescent beads, fixed tissue, and *in vitro* GCaMP6s brain slice imaging. An electrically tunable lens (Optotune, Dietikon, Switzerland) attached to the back of the objective was used for fast focus adjustments in the fluorescence imaging experiments but not for the beam characterization. The fluorescent beads, fixed tissue, and *in vitro* imaging used 200 ms, 500 ms, and 100 ms camera exposure times, respectively. The packaged probe was attached to a 4-axis micro-manipulator for positioning the probe in the characterization and imaging experiments. The shanks were aligned to the insertion axis of the micro-manipulator, and in the imaging experiments, the insertion angle was selected for sheet-normal imaging. Since the fiber bundle was not polarization-maintaining and the probe was polarization-sensitive, the bundle was fixed in position during imaging experiments to minimize polarization fluctuations.

### Animals

All experimental procedures described here were reviewed and approved by the animal care committees of the University Health Network in accordance with the guidelines of the Canadian Council on Animal Care. Adult Thy1-GCaMP6s mice [2] (The Jackson Laboratory, Bar Harbor, ME, USA, stock number 025776) and C57BL/6 mice (Charles River Laboratories, Wilmington, MA, USA) were kept in a vivarium maintained at 22°C with 12-h light on/off cycle. Food and water were available ad libitum.

### Fixed tissue preparation

Fixed tissue was prepared from a Thy1-GCaMP6s mouse, postnatal day 172, as ≈ 1 mm thick transverse slices from the hippocampus (dentate gyrus). Briefly, the animal was anesthetized via an intra-peritoneal injection of sodium pentobarbital (75 mg/kg, Somnotol, WTC Pharmaceuticals, Cambridge, Ontario, Canada) and transcardially perfused with 1× phosphate-buffered saline (PBS) followed with paraformaldehyde (PFA) (4%). Then the extracted brain was kept in PFA at 4°C for 12 hours. After fixation, the hippocampal slices were prepared in 1× PBS with a vibratome (VT1200S, Leica Biosystems, Wetzlar, Germany).

### *in vitro* imaging brain slice preparation

Brain slices were prepared from 30-60 day old Thy1-GCaMP6s and C57BL/6 mice for the *in vitro* GCaMP6s and calcium dye imaging experiments, respectively. The animals were anesthetized with an intra-peritoneal injection of sodium pentobarbital (75 mg/kg) and transcardially perfused with cold (4°C) N-methyl-D-glucamine (NMDG) recovery solution [45] prior to decapitation. The brain was quickly dissected, brain tissues were glued on a vibratome stage, and 450 *μ*m thick slices were prepared with the vibratome using iced NMDG solution. The brain slices were then stabilized in NMDG solution at 34°C for 12 min while being aerated with carbogen (95% O_2_, 5% CO_2_). Only for experiments with the Cal-520 AM calcium dye, following a 12-minute recovery period, slices were rinsed and then bathed in a Cal-520 AM solution (AAT Bioquest, Sunnyvale, CA, USA) for 60 - 90 minutes at 37°C. For all *in vitro* experiments, the slices were then maintained in room temperature incubation solution [45] for 1-8 hours prior to imaging. The Cal-520 AM solution was aerated with carbogen and consisted of 50 *μ*g of Cal-520 AM mixed with 20 *μ*L of 20% Pluronic F-127 in dimethyl sulfoxide (DMSO) (Sigma-Aldrich, St. Louis, MO, USA) and then diluted in 4 - 6 mL of incubation solution to a final concentration of 7 - 10 *μ*Mol. During imaging of a slice, the slice was mounted in a perfusion chamber with a constant flow of rodent artificial cerebrospinal fluid (aCSF) solution [45] aerated with carbogen. A 100 - 200 *μ*Mol solution of 4-aminopyridine (4-AP) was added to the aCSF bath to put the neurons in a hyperexcitable state for increased neuronal activity [37]. For the Thy1-GCaMP6s imaging, a transverse slice prominently featuring the auditory cortex was chosen, and for the Cal-520 AM imaging, a sagittal slice prominently featuring the cerebellum was chosen. The cerebellum was chosen since we observed that it had high neuron activity density. The cerebellum could not be chosen for the Thy1-GCaMP6s experiment due to the low labeling density in the cerebellum for this strain [2].

## V. Appendix

Four supplementary videos are included in this work, and descriptions are below.

**Video S1:** Top-down microscope imaging of Probe 3 with switching between light sheets. The video is real-time.

**Video S2:** Side microscope fluorescence imaging of Probe 1 immersed in a fluorescein solution with switching between the 5 light sheets. The power of the sheets was roughly equalized here by not optimally aligning the MEMS mirror for the sheets with relatively high transmission. Additional ambient illumination was applied to make the shanks visible. The video is real-time.

**Video S3:** Fluorescence imaging of 3 μm diameter fluorescent beads suspended in an agarose block with light-sheet illumination from Probe 1 and imaging using the epifluorescence microscope above the sample. At about 17 s, the illumination is switched from light-sheet probe illumination to epi-illumination from the microscope. During light-sheet probe illumination, switching between Sheets 1, 3, and 5 is performed. The tunable lens is synchronized to the sheet switching to focus the collection optics on the depth planes corresponding to each sheet. After switching to epi-illumination, the tunable lens focus switching continues and shows epifluorescence imaging of the same depth planes imaged with light-sheet probe illumination. The video is real-time.

**Video S4:** *In vitro* calcium imaging with epi-illumination of a Cal-520 AM loaded brain slice from a wild type mouse. This is a sample of the calcium imaging video corresponding to Fig. S8 in the Supplementary Materials. The video is from the same experiment as Video 3 but with epi-illumination instead of light-sheet probe illumination. The video is real-time.

## Acknowledgments

This work was supported by NIH awards NS090596 and NS099726, Canadian Institute of Health Research award FRN151949, and the Natural Sciences and Engineering Research Council of Canada award CHRPJ 508406. Funding support from the Canadian Foundation for Innovation and Ontario Research Fund is also gratefully acknowledged. W.D.S. was supported by the Kavli Nanoscience Institute Prize Postdoctoral Fellowship in Applied Physics and Materials Science. A.F. is supported by the Clinician Investigator Program - University of Manitoba. The authors thank Michael Chang and Azadeh Naderian at the Krembil Research Institute for their assistance with the animal colonies and genotyping. The authors also thank Alex Jacob in the group of Professor Sheena Josselyn at SickKids Research Institute (Toronto, Canada) for his advice on GCaMP6 functional imaging.

## Data availability

The data are available from the corresponding authors upon reasonable request.

## Additional information

Supplementary information is available for this paper.

