## Supplementary Materials for "Implantable photonic neural probes for light-sheet fluorescence brain imaging"

[†] *Equal contribution*

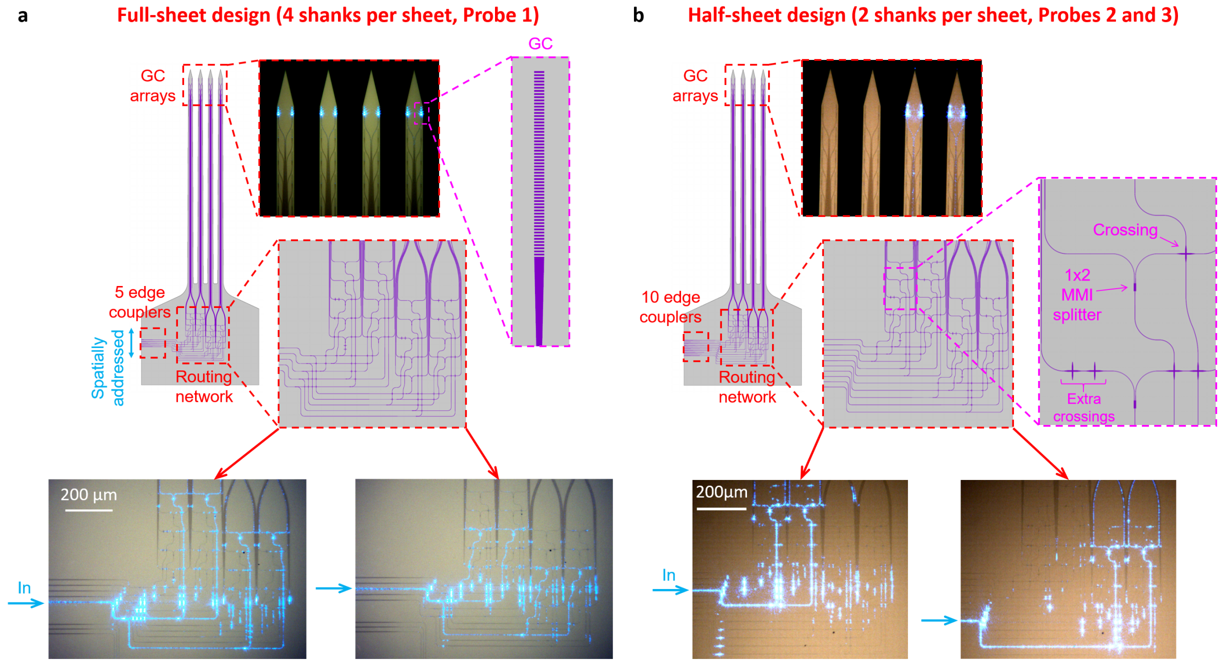


**Figure S1: Additional details of the light-sheet neural probe designs.** Comparison of the light-sheet neural probe designs with (a) 4 shanks per sheet and (b) 2 shanks per sheet. The design in (a) has 5 independent sheets and 5 edge couplers, and the design in (b) has 10 independent sheets and 10 edge couplers. Schematics of the probes with magnified views of the routing networks are shown. Microscope images of the grating couplers (GCs) emitting a single sheet are shown for both designs. Microscope images of the routing networks with optical inputs to 2 different edge couplers are also shown for both designs. The rightmost inset in (b) shows the $1 \times 2$ MMI power splitters and waveguide crossings used in the routing networks. Our design and characterization of these devices is described in [34]. For transverse-electric (TE) polarized light at a wavelength of 488 nm, the measured excess loss of the $1 \times 2$ MMI power splitters was $\approx 0.5$ dB per power splitter, and the loss of the crossings was $< 0.2$ dB/crossing [34].

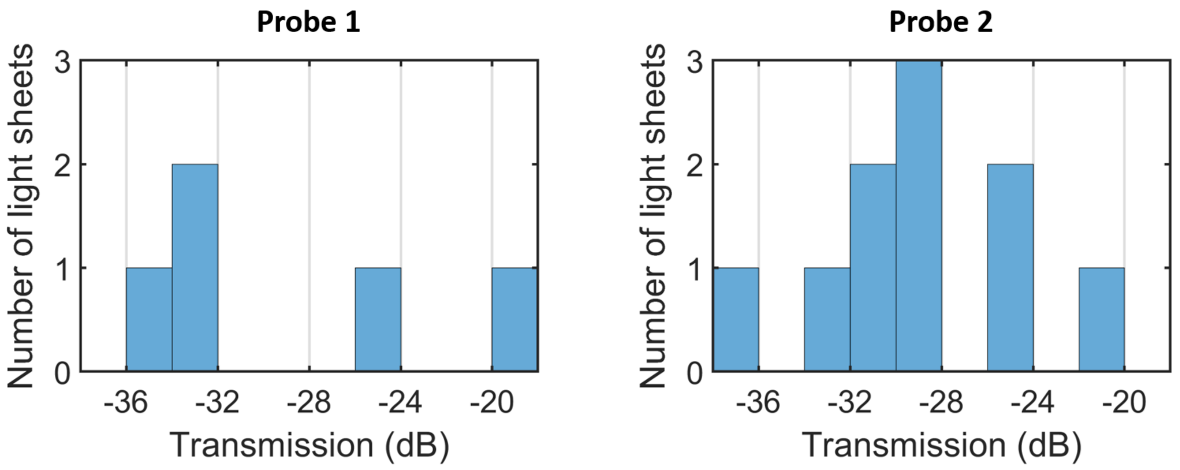


**Figure S2: Histograms of the optical transmissions of the light sheets from packaged Probe 1 and Probe 2.** The transmission is the ratio of the emitted power from the GCs to the input laser power to the scanning system.

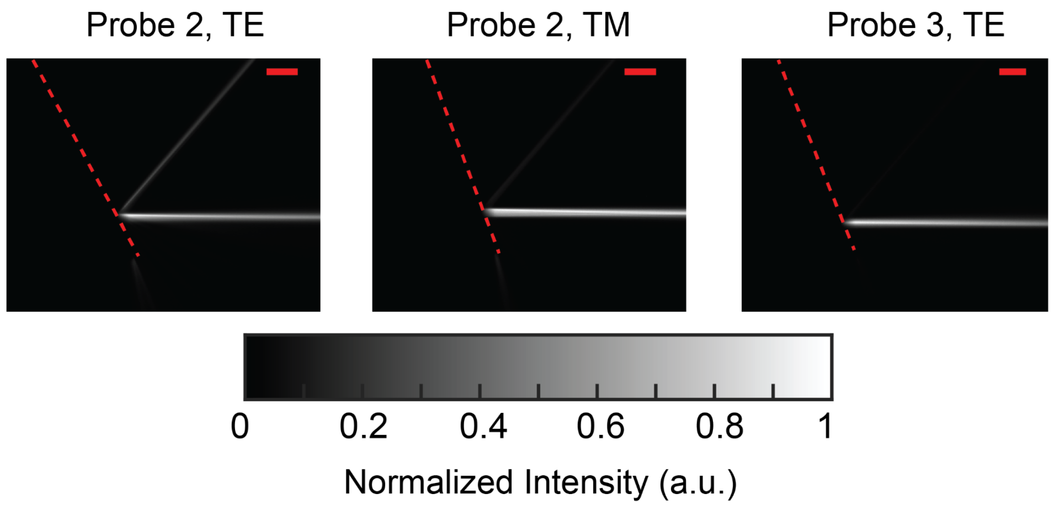


**Figure S3: Additional fluorescence images of side sheet profiles from light-sheet neural probes immersed in fluorescein.** Left: Probe 2 for the TE polarization. Center: Probe 2 for the TM polarization. Right: Probe 3 for the TE polarization. The dashed red lines delineate the top surface of the shanks. The scale bars are 100 µm.

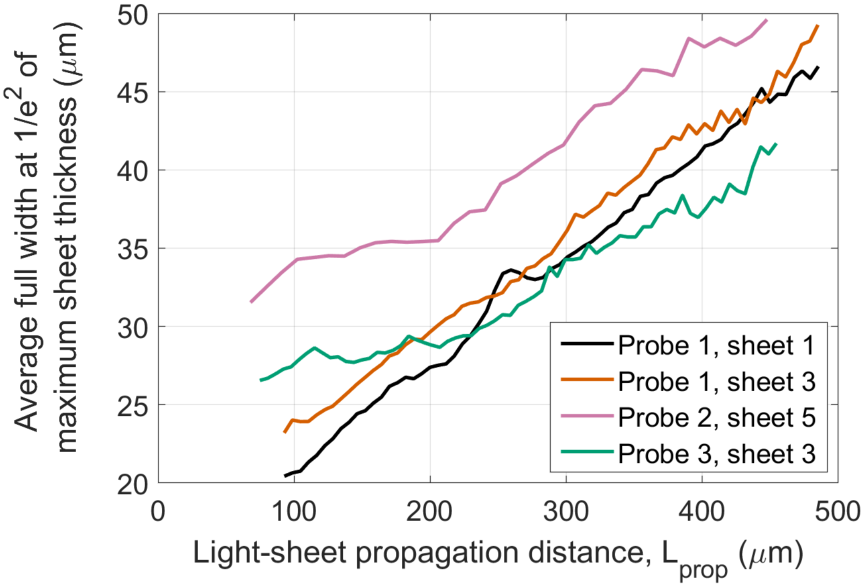


**Figure S4: Additional data for the free-space light-sheet profile measurements in Fig. 4.** Average full width at $1/e^2$ of maximum sheet thickness versus light-sheet propagation distance for sheets from Probes 1 – 3. The sheet thicknesses are averaged over the sheet width, the vertical axis in Fig. 4(c). The FWHM sheet thickness for Probe 1 – Sheet 2 is shown in Fig. 4(b), but the $1/e^2$ sheet thickness could not be measured accurately and is not shown here because the relatively weak transmission of the sheet resulted in $1/e^2$ intensity levels close to the noise floor of the measurement. In general, the $1/e^2$ sheet thicknesses increase linearly with propagation distance, while the Probes 2 and 3 FWHM sheet thicknesses in Fig. 4(b) are nonlinear with propagation distance, a result of the evolution of the sheet profiles with propagation distance.

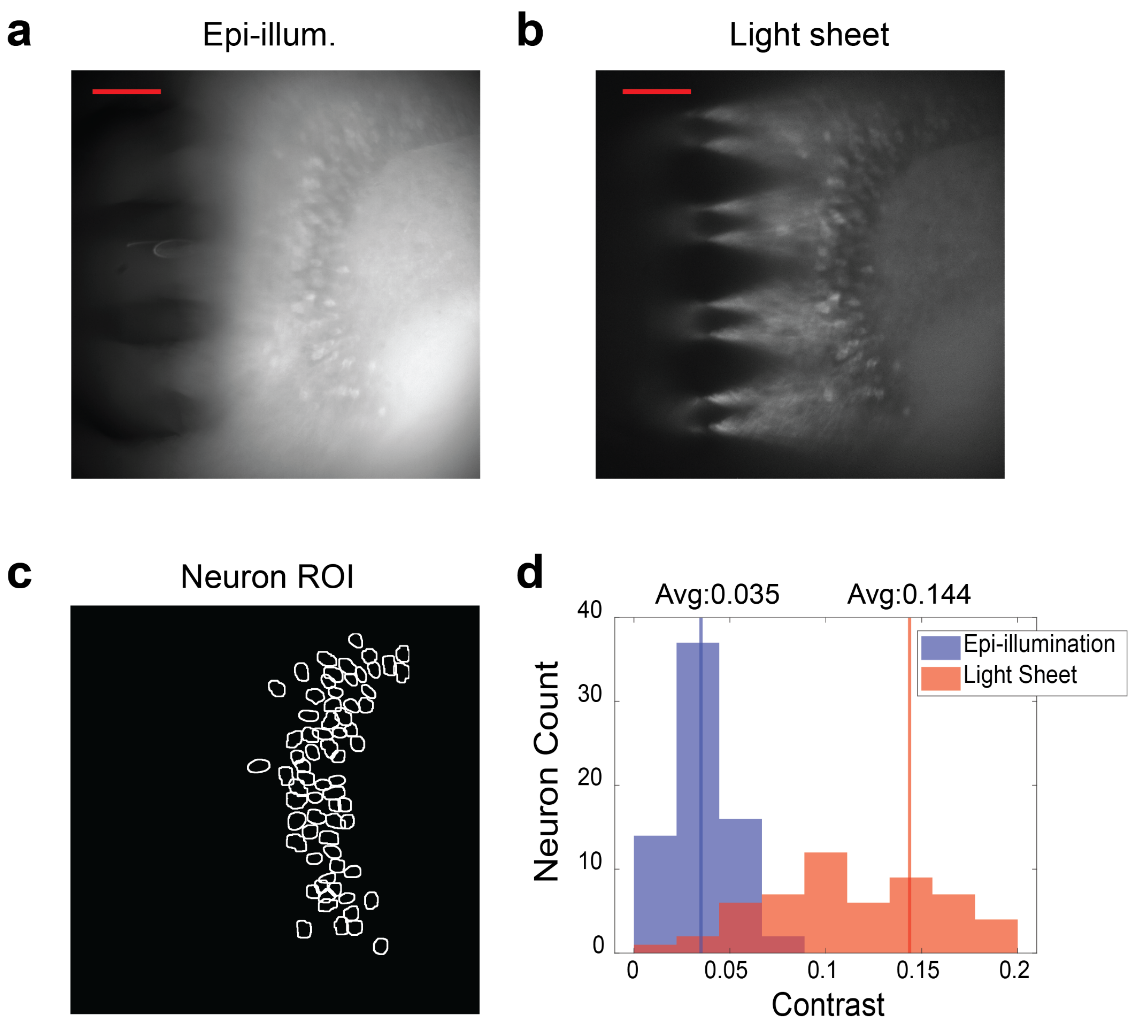


**Figure S5: Additional data for the Thy1-GCaMP6s fixed mouse brain tissue imaging in Fig. 6(a).** Microscope epi-illumination and light-sheet neural probe illumination (Sheet 3, Probe 1) are compared using a reduced light-sheet intensity relative to Fig. 6(a). (a-b) Top-down fluorescence images of (a) epi-illumination and (b) light-sheet illumination. The epi-illumination image data in (a) is the same as in Fig. 6(a). The scale bars are 100 µm. (c) Regions of interest (ROIs) of identified neurons. (d) Histograms of image contrast for the identified neurons. The light-sheet image was taken during the turn-off transient of the shutter gating the input laser beam to the scanning system, and therefore, the light-sheet image is an average of multiple intensities over the camera exposure time (500 ms). The average intensity of the light-sheet image is 2.85× lower than that of Fig. 6(a). The light-sheet average neuron contrast in (d) was not significantly changed compared to the higher sheet intensity image in Fig. 6(a).

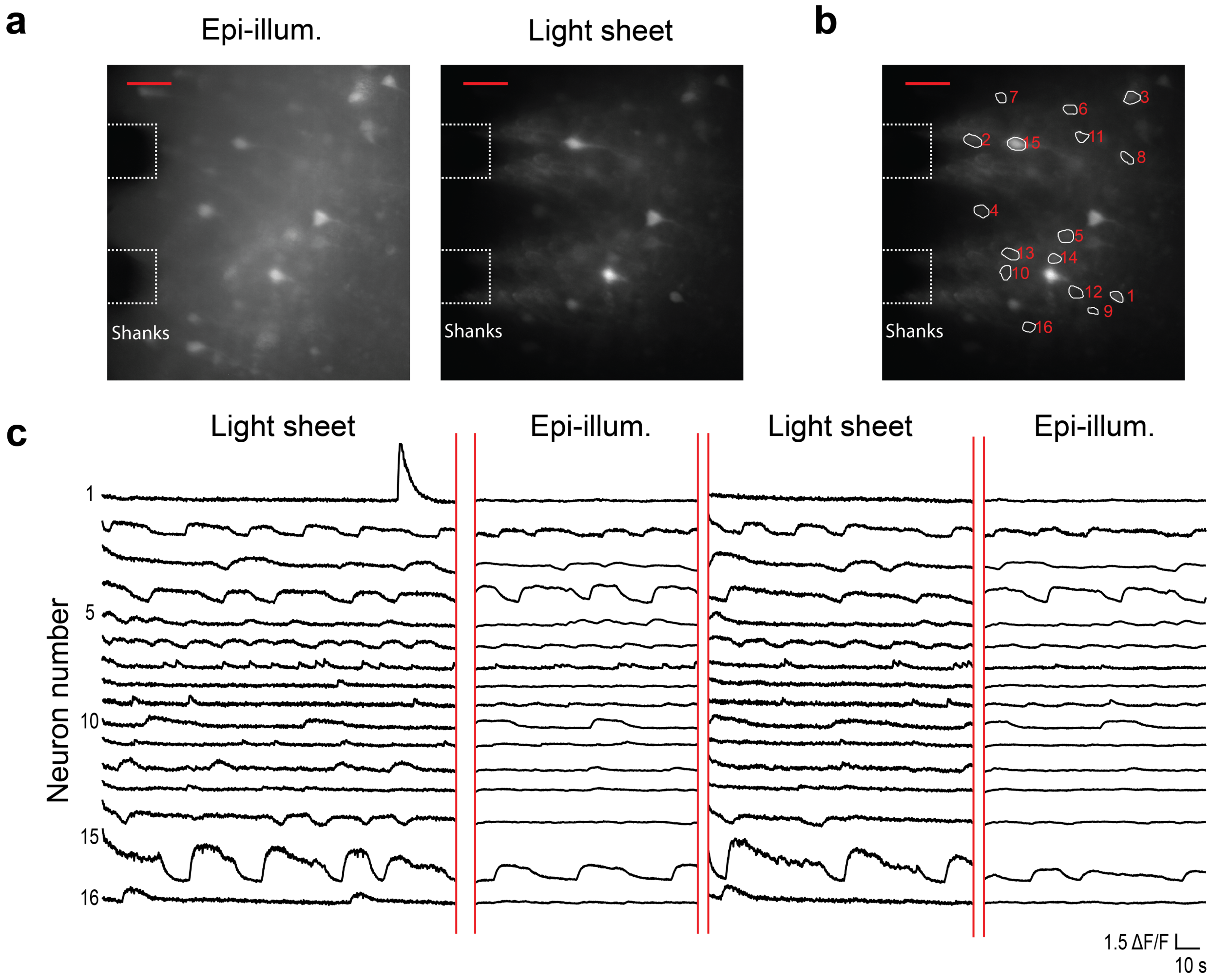


**Figure S6: Additional data for the *in vitro* Thy1-GCaMP6s mouse brain slice calcium imaging in Fig. 7 showing the complete time traces as the illumination was switched between light-sheet and epi-illumination.** (a) Maximum projection images of the first segments of the recorded video for light-sheet probe illumination and microscope epi-illumination. The scale bars are 50 µm. (b) Light-sheet maximum projection image from (a) with regions of interest (ROIs) for identified neurons shown. (a-b) are repeated from Fig. 7 to show the neuron ROIs for the time traces in (c). (c) Fluorescence change, ΔF/F, time traces of the neurons identified in (b) as the illumination was switched between light-sheet and epi-illumination. Gaps in the time traces correspond to periods when no illumination was applied. The time traces correspond to the video used to generate the neuron contrast box plots in Fig. 7(d).

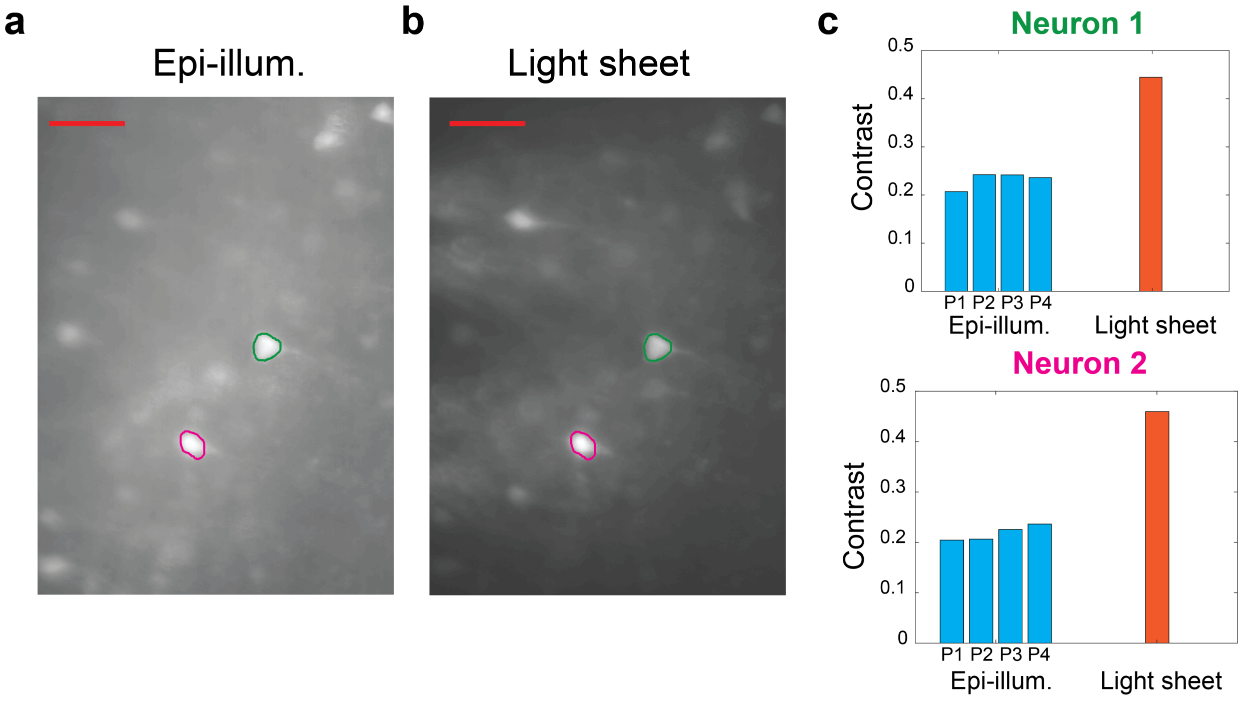


**Figure S7: Image contrast comparison of 2 neurons in the Thy1-GCaMP6s brain slice imaged in Fig. 7 at multiple microscope epi-illumination intensities and with a single light-sheet probe illumination intensity (Sheet 5, Probe 2).** (a-b) Brain slice fluorescence images with (a) epi-illumination and (b) light-sheet probe illumination; the ROIs of the 2 neurons are indicated, and the scale bars are 50 µm. (c) Image contrast of the neurons at 4 epi-illumination intensities, P1-P4, and with light-sheet illumination using the intensity in Fig. 7. The estimated epi-illumination intensities (P1, P2, P3, and P4) were 0.7, 1.1, 1.9, and 2.8 $mW/mm^2$, respectively, and the corresponding contrasts span 20.7% to 24.2% for neuron 1 and 20.5% to 23.6% for neuron 2. The epi-illumination neuron contrasts are consistently lower than the light-sheet contrasts. The neuron image contrast is calculated using Eq. 1 in Supplementary Note 1, averaging over 50-107 s of video. The 2 neurons were selected for their ease of identification in both epi- and light-sheet illumination images. At each illumination condition, over the time averaging window, the time-varying neuron intensity averaged over the ROI had a standard deviation relative to the median of 2.7 – 4.5% for epi-illumination and 11.9 – 12.8% for light-sheet illumination.

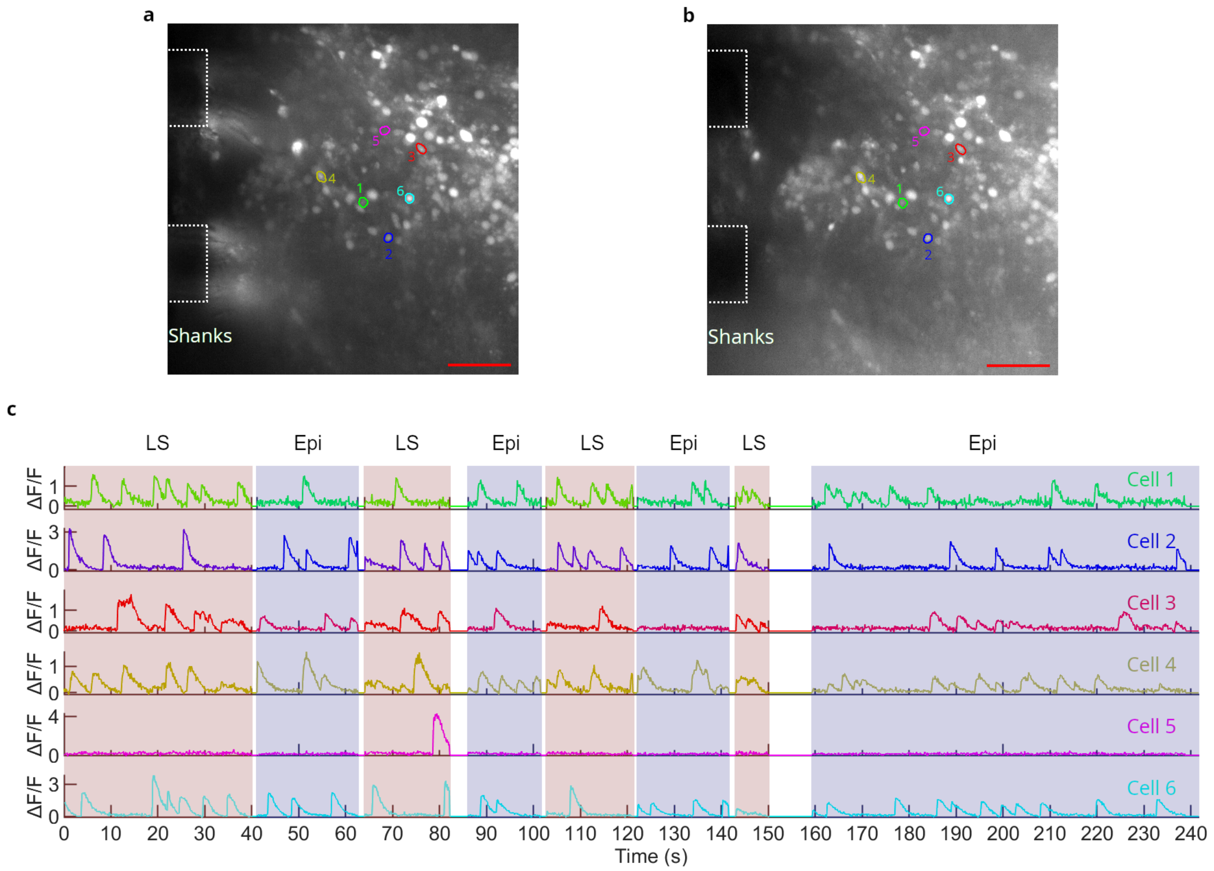


**Figure S8: Additional data for the *in vitro* Cal-520 AM loaded mouse brain slice calcium imaging in Fig. 8.** (a-b) Maximum projection fluorescence images of segments of the recorded video for (a) light-sheet and (b) epi-illumination. Annotations show the approximate positions of the shanks in the image plane, and the scale bars are 50 µm. The ROIs of the 6 cells selected for time traces in Fig. 8 are shown. (c) ΔF/F time traces of the 6 cells identified in (a-b) as the illumination was switched between light-sheet ("LS") and epi-illumination ("Epi"). The colors and numbers of the cell ROIs correspond to the time traces. Gaps in the time traces correspond to periods when no illumination was applied. The maximum projection images in (a-b) are taken over the first light-sheet and epi-illumination time segments in (c).

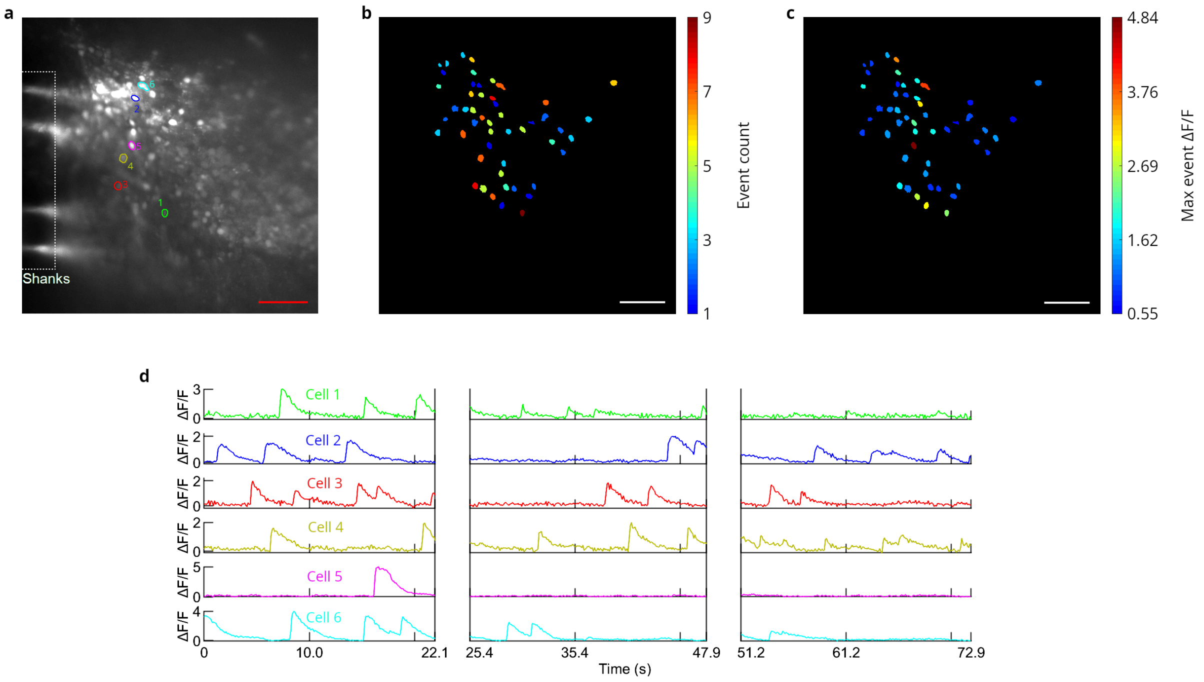


**Figure S9: Additional *in vitro* calcium imaging of a Cal-520 AM loaded mouse brain slice using light-sheet probe illumination from Probe 2.** The brain slice is the same as Figs. 8 and S8, but the probe was inserted at a different position in the slice. (a) Maximum projection fluorescence image of a segment of the recorded video with annotations to show the approximate position of the shanks in the image plane. (b) Maximum observed ΔF/F and (c) number of calcium events observed for all identified cells. (b-c) are of the same scale as (a), and the scale bars are 50 µm. (d) ΔF/F time traces of 6 cells. The number and color-coded ROIs for these cells are shown in (a). Breaks in the time traces correspond to periods when the illumination was switched to epi-illumination. The maximum projection image in (a) is taken over the first time segment in (d).

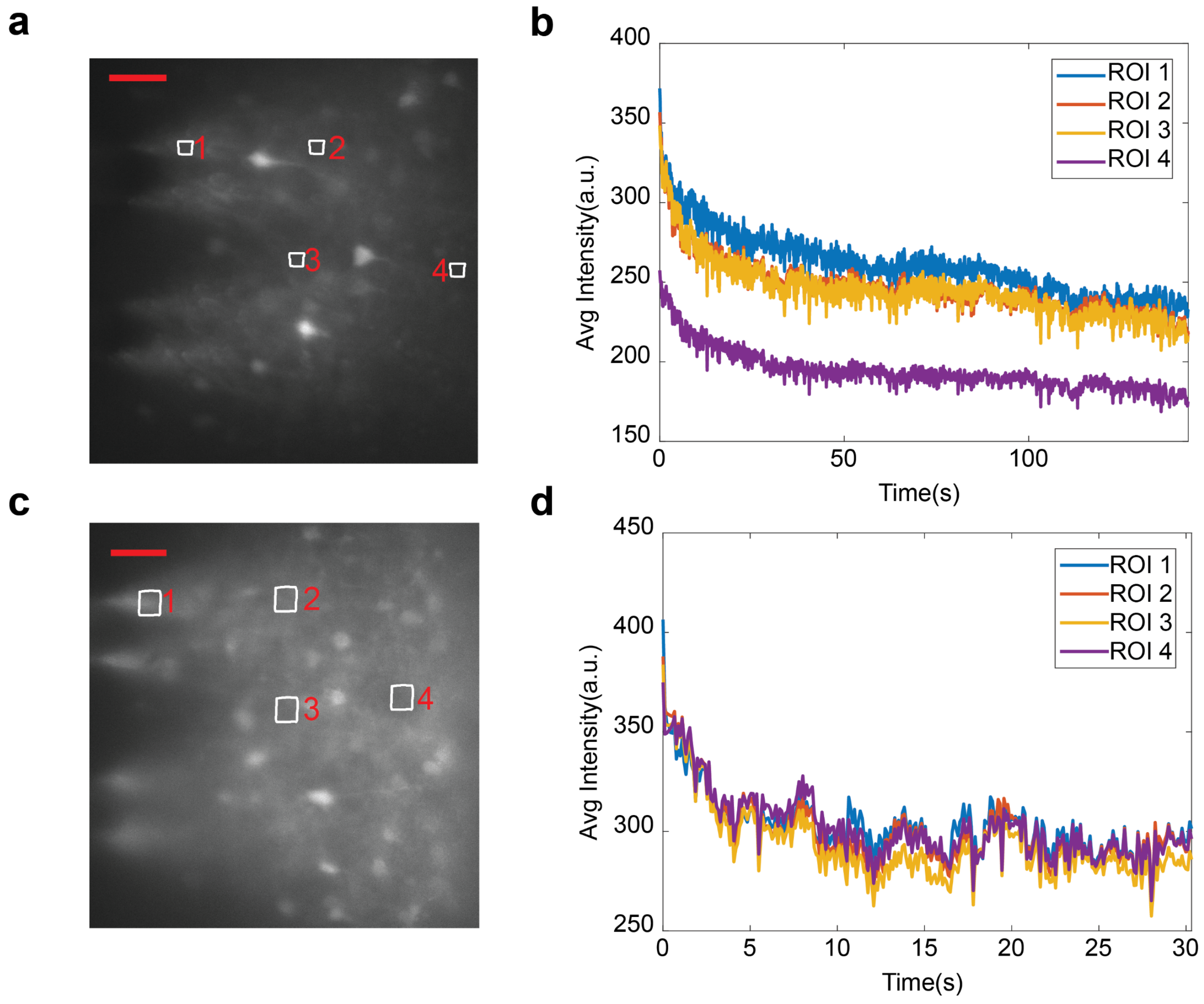


**Figure S10: Example light-sheet neural probe (Probe 2) fluorescence image intensity turn-on transients for *in vitro* Thy1-GCaMP6s mouse brain slice calcium imaging.** (a,c) Fluorescence images of 2 brain slices from the same mouse with 4 ROIs labeled in each. The ROIs are outside the observable neurons and were selected to sample the neuropil fluorescence at different positions in the sheet. (b,d) Image intensity time traces averaged over each ROI; (b) corresponds to the imaging in (a), and (d) corresponds to (c). The beginning of the time-axis corresponds to the time at which the light-sheet illumination was turned on. The scale bars in (a) and (c) are 50 µm. The data in (a) and (b) are part of the same imaging experiment performed in Fig. 7. The turn-on transient in (b) and (d) may be a combination of thermal expansion of the probe packaging upon absorbing stray input light not coupled to the chip and photobleaching of the brain slice.

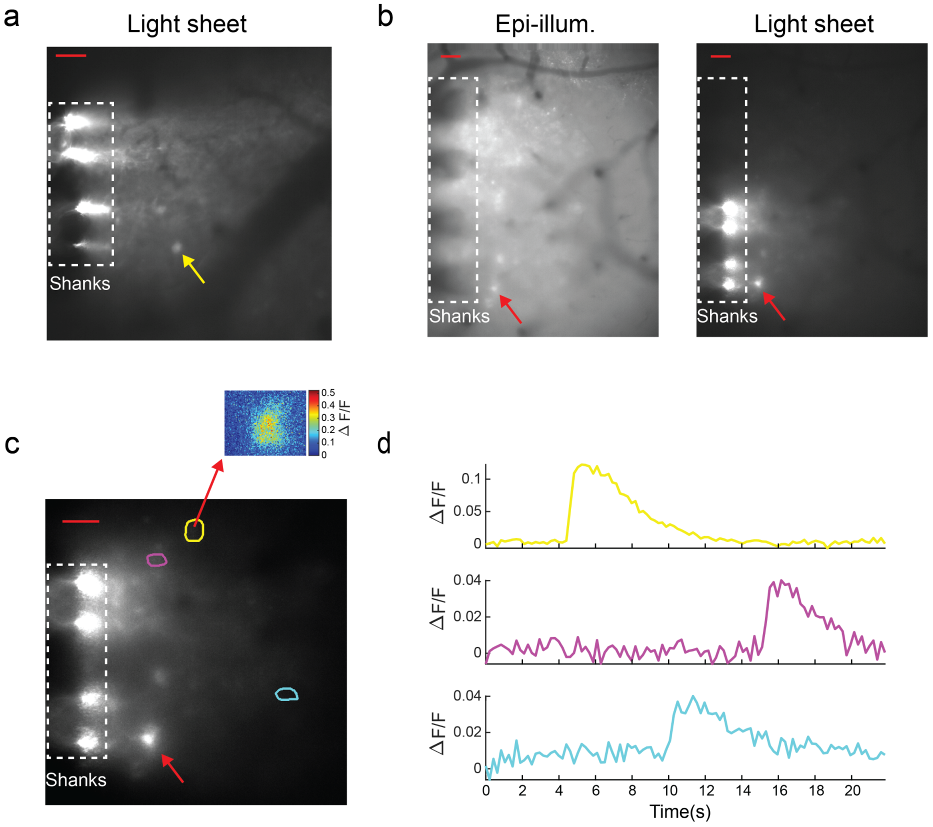


**Figure S11: *In vivo* fluorescence imaging of the parieto-temporal lobe at the approximate location of the somatosensory cortex of an anesthetized Thy1-GCaMP6s mouse.** (a) Fluorescence image with light-sheet probe illumination. (b) Light-sheet and epi-illumination fluorescence images with the probe inserted into the cortex at a different position relative to (a). (c) Maximum projection fluorescence image over a segment of the video corresponding to (b) with light-sheet illumination. The image is annotated with the ROIs of 3 neurons identified by fluorescence changes consistent with GCaMP6s. The inset shows the fluorescence change, ΔF/F, at the peak of an observed calcium event for a rectangular region around and including one of the ROIs. (d) ΔF/F time traces of the 3 neurons; the colors of the traces are matched to the colors of the ROIs in (c). The neuron corresponding to the red arrow in (b-c) exhibited higher contrast for light-sheet relative to epi-illumination by a factor of ≈ 3.4×. The fluorescent point source corresponding to the yellow arrow in (a) could not be identified as a neuron due to a lack of time-dependent fluorescence; a contrast of enhancement of ≈ 2.7× was observed for light-sheet illumination compared to epi-illumination. The scale bars of (a-c) are 50 µm, and the dashed boxes denote the approximate positions of the shanks in the image planes. The intensity grayscales in (a-b) are set by the maximum and minimum pixel values outside the dashed boxes, and the grayscale of (c) is set by 99$^{th}$ and 1$^{st}$ percentiles of pixel intensities. The calculation of ΔF/F here follows the calculation in Supplementary Note 3 but without background subtraction in the baseline fluorescence; ΔF/F in (c) also uses a 5 frame moving average and the annular ROI for background calculations is 14 µm wide. We previously reported this data in [31].

### Supplementary Note 1: Fixed tissue imaging data analysis

Neuron regions of interest (ROIs) were identified within the fixed tissue fluorescence images using the following algorithm. First, background subtraction was performed on the images. The background was estimated with a moving average window across the image. Next, adaptive thresholding followed by a morphology operation was performed on the image to extract regions with prominent peaks [46]. Local peak detection with watershed segmentation was then applied to each extracted neuron ROI for separating ROIs merged together due to close proximity. Manual intervention was done to reject false detection and to include undetected neurons. Neuron ROIs were defined manually using Fiji [47] to approximate closely the outlines of the neuron somas. In Fig. 6(a), 47 neuron ROIs were defined by the algorithm above and 22 neuron ROIs were defined manually. In Fig. 6(b), all ROIs were defined manually.

For the contrast calculation, dark-frame subtraction was performed for ambient illumination removal. The ambient level in the frame was estimated by averaging over 35 frames for Fig. 6(a) and 44 frames for Fig. 6(b) with the absence of excitation illumination. Then, each neuron mask was dilated by 5.7 $\mu$m (20 pixels) to form an annular region surrounding the neuron ROI. Next, we calculated the contrast of each neuron with the following equation:

$$contrast = \frac{F_{neuron} - F_{background}}{F_{background}} \tag{1}$$

Here, $F_{neuron}$ is the mean fluorescence value within the ROI of the neuron mask and $F_{background}$ is the mean fluorescence level within the annular mask. Portions of the annular masks overlapping the ROIs of other neurons were excluded from the background calculation.

For the fixed tissue and *in vitro* imaging, image and statistical analyses and plotting were performed using MATLAB (MathWorks Inc., Natick, MA, USA) and Python (version 3.6.7) [48]. The maximum and minimum of the intensity scales for the grayscale fixed tissue fluorescence images in Fig. 6 were set as the maximum and minimum of all pixel values in the corresponding raw images.

**Supplementary Note 2: Light-sheet and epi-illumination intensity estimates**

For the fixed tissue fluorescence imaging in Fig. 6, the epi-illumination intensity at the surface of the brain slice was measured to be approximately 0.5 $mW/mm^2$. The Sheets 3 and 4 output powers from the neural probe were approximately 60 μW and 3 μW, respectively. Neglecting tissue scattering, at a propagation distance of 200 μm from the probe, assuming the sheet width and thickness from Figs. 3(b) and 4(b), respectively, yields Sheets 3 and 4 intensity estimates of ≈11 $mW/mm^2$ and 0.6 $mW/mm^2$, respectively.

For the GCaMP6s and Cal-520 AM *in vitro* imaging in Figs. 7, 8, S6, and S8, the epi-illumination intensities were measured to be approximately 0.3 $mW/mm^2$ and 1.0 $mW/mm^2$, respectively. For light-sheet illumination, neglecting tissue scattering and assuming the sheet width and thickness from Figs. 3(c) and 4(b), at a propagation distance of 200 μm from the probe, the light sheet intensity estimate in these experiments is 13 $mW/mm^2$.

Tissue scattering is expected to significantly reduce the intensity of the light-sheet and epi-illumination. Rather than quantifying this effect and matching the intensities inside the tissue for comparing light-sheet and epi-illumination, we have varied the illumination intensity and confirmed no large changes in the contrast enhancement of light-sheet probe illumination relative to epi-illumination. Figure S5 shows fixed tissue imaging with reduced light-sheet illumination intensity (relative to Fig. 6) and no significant degradation in the ratio of light-sheet to epi-illumination image contrast. In Fig. S7, the epi-illumination intensity is varied over a ≈ 4× range for the Thy1-GCaMP6s mouse brain slice imaged in Fig. 7. For the two neurons analyzed, the image contrast does not continuously increase with epi-illumination intensity and remains significantly less than that of light-sheet illumination in all cases.

### Supplementary Note 3: *In vitro* imaging data analysis

First, any vibration in the calcium imaging video was corrected with the rigid motion correction method. Then, active neurons were detected using the constrained non-negative matrix factorization (CNMF) algorithm. Neurons with SNR below 2 and spatial correlation of the footprint during active phase below 0.85 were filtered to reduce the false detection rate. The neuron detection described above was performed with the analysis code provided in [49]. Next, manual inspection of the video was conducted to ensure the quality of the detected signals. For the GCaMP6s slice analysis, manual inspection of the time-dependent fluorescence traces was used to remove falsely detected neurons and to include undetected neurons; neuron ROIs were defined manually using Fiji. For the Cal-520 AM slice analysis, neuron ROIs were defined by the analysis code in [49], and manual inspection of the fluorescence change, $\Delta F/F$, traces was used to remove detected neurons with any of the following: 1) no detected peaks in $\Delta F/F > 0.5$ (using the Matlab *findpeaks* function), 2) $\Delta F/F$ noise levels high enough to be detected as peaks using the *findpeaks* function, 3) $\Delta F/F$ traces without characteristics consistent with Cal-520 AM calcium event dynamics.

$\Delta F/F$ traces were calculated as follows. For each identified active neuron, a binary mask was generated, and an annular region with a width of 6 $\mu$m surrounding the neuron ROI was also generated. The fluorescence of each neuron was spatially averaged over the neuron ROI, resulting in a time-dependent trace $F_{raw}$ for each neuron. To partially correct for neuropil fluorescence and illumination fluctuations, background removal was performed on each $F_{raw}$ with the background signal ($F_{bg}$) estimated by the first percentile of the fluorescence intensity within the annular region surrounding the neuron ROI [50, 51]. $F_{baseline}$ is the baseline of $F_{raw} - F_{bg}$, and for the Cal-520 AM slice analysis, $F_{baseline}$ was determined by the average of the lowest 10% of the values within a 7 s wide moving window swept over $F_{raw} - F_{bg}$ [52]. Moving windowing was not used for GCaMP6s slice data due to the long duration of some of the calcium events. Instead, $F_{baseline}$ was taken as the average of the lowest 10% of the whole $F_{raw} - F_{bg}$ time trace for a video segment (segments defined in Fig. S6). The fluorescence change, $\Delta F/F$, for each neuron was then calculated as:

$$\frac{\Delta F}{F} = \frac{F_{raw} - F_{bg} - F_{baseline}}{F_{baseline}} \quad (2)$$

A small number of ROIs with dimensions significantly smaller than the surrounding cells are present in the *in vitro* Cal-520 AM imaging analysis in Figs. 8(b) and (c). Such an ROI is an "island" of a larger adjacent ROI, both belonging to the same cell. These islands occur because the CNMF algorithm does not guarantee an enclosed shape for a neuron ROI [49]. The islands were not considered to be separate from their parent ROIs for the calculations. These islands did not occur in the analysis of the *in vitro* GCaMP6s brain slice imaging in Fig. 7 since the ROIs were defined manually.

The maximum and minimum of the intensity scale of each grayscale *in vitro* GCaMP6s brain slice image in Figs. 7, S6, and S7 were set as the maximum and minimum pixel intensity values of the raw fluorescence (or maximum projection fluorescence) image (using MATLAB). The *in vitro* Cal-520 AM grayscale maximum projection fluorescence images in Figs. 8, S8, and S9 were generated from raw video data using Fiji with the grayscales set using the *autoscale* function.

Videos 1 – 3 and S1 – S4 were processed using Fiji. To generate Videos 1 – 3 and S4, the raw 16-bit calcium imaging videos were cropped to the areas analyzed in Figs. 7 and 8, the grayscales (brightness and contrast) were adjusted, and the videos were then converted to mp4 video files (8-bit, h.264 compression). The procedure for setting the intensity grayscales of Videos 1 – 3 and S4 is as follows. For each video, the maximum pixel intensity of each frame was compiled into a distribution of maxima and the 99$^{th}$ percentile of the distribution was found. For each video, the 1$^{st}$ percentile of the distribution of minimum pixel intensities of all frames was also found. Due to the similarity of the maximum fluorescence intensities in the GCaMP6s imaging in Videos 1 and 2, the same intensity grayscale was applied to both videos; the maximum and minimum of the grayscale were set as the larger 99$^{th}$ percentile of maxima and the smaller 1$^{st}$ percentile of minima, respectively, of the two videos. The maximum fluorescence values for the Cal-520 AM imaging in Videos 3 and S4 differed significantly and required different intensity grayscales for each video. The maximum and minimum of each grayscale in Videos 3 and S4 were set as the 99$^{th}$ and 1$^{st}$ percentiles of the distributions of maxima and minima of each video, respectively.

### Supplementary Note 4: *In vivo* imaging methods

All experimental procedures described here were reviewed and approved by the animal care committees of the University Health Network in accordance with the guidelines of the Canadian Council on Animal Care. First, the Thy1-GCaMP6s mouse (postnatal day 47) was anesthetized, by induction with 5% and maintenance with 1% to 2% isoflurane/oxygen anesthetic, and secured in a stereotaxic frame (Model 902, David Kopf Instrument, Tujunga, CA, USA). The mouse head was secured with ear bars in the stereotaxic frame. A craniotomy was performed over the parieto-temporal lobe with an electric drill by removing a square portion of the skull posterior to the bregma and anterior to the lambda bony landmarks (1 mm lateral to midline and 1 mm medial to the superior attachment of the temporalis muscle). The dura was gently removed with a 30 gauge needle for higher image quality and easier probe insertion. Once the cranial opening was complete, the stereotaxic frame – with the mouse secured – was moved under the epifluorescence microscope and the brain surface was irrigated with saline. Initial epifluorescence brain imaging was performed to identify the somatosensory cortex, and an area relatively free of cortical blood vessels in the parieto-temporal lobe at the approximate location of the somatosensory cortex was chosen for probe insertion. Next, the probe was slowly inserted into the brain to a maximum depth of 200 µm via the micro-manipulator, and then, fluorescence imaging was performed using both probe- and epi-illumination. The probe insertion angle was set so that the light sheet was roughly parallel to the surface of the brain.

The *in vivo* fluorescence imaging used a 200 ms exposure time and 2×2 binning; binning was not used in the other imaging experiments in this work. The *in vivo* fluorescence imaging results are summarized in Fig. S11.

### Supplementary Note 5: Additional details of the neural probe used for *in vivo* imaging

The light-sheet neural probe used for the *in vivo* fluorescence imaging in Fig. S11 was a prototype that preceded our foundry fabricated probes. This probe was fabricated using the method described in [26] at the California Institute of Technology's Kavli Nanoscience Institute cleanrooms. Briefly, these probes were fabricated on 100 mm diameter silicon-on-insulator (SOI) wafers with a 15 μm thick Si device layer and a 2 μm thick buried oxide layer. A 1.5 μm thick thermally grown $SiO_2$ layer was used as the bottom cladding of the waveguides. A 200 nm thick silicon nitride (SiN) waveguide layer deposited by low pressure chemical vapor deposition (LPCVD) was patterned and etched by electron beam lithography and an inductively coupled plasma (ICP) pseudo-Bosch etch process. The waveguide top cladding was a 1 μm thick plasma enhanced chemical vapor deposition (PECVD) $SiO_2$ layer. Front- and back-side deep reactive ion etching (DRIE) was used to define the probe shape, edge coupler facets, and selectively thin the shanks to a thickness of $\approx$ 18 μm, while leaving the base of the probe chips (containing the routing network and edge couplers) at the full $\approx$ 300 μm thickness of the SOI wafer.

Other than the thickness, the designed dimensions of these neural probes were nearly identical to those of the foundry fabricated neural probes. The shank pitch was 135 μm, the shank length was 3 mm, the pitch of the rows of grating couplers (GCs) for each sheet was 80 μm, the GC socket width was 1 μm, and the GC period was 400 nm with a 50% nominal duty cycle.
